# A signalling rheostat controls chromosome segregation fidelity during early lineage specification and neurogenesis by modulating DNA replication stress

**DOI:** 10.1101/2023.07.18.549463

**Authors:** A De Jaime-Soguero, J Hattemer, A Haas, A Bufe, B Di Marco, N Böhly, JJM Landry, B Schoell, VS Rosa, L Villacorta, Y Baskan, S Androulaki, M Trapp, V Benes, B Das, M Shahbazi, A Jauch, U Engel, A Patrizi, R Sotillo, J Bageritz, J Alfonso, H Bastians, SP Acebrón

**Affiliations:** Centre for Organismal Studies (COS), Heidelberg University, D-69120 Heidelberg, Germany; Georg-August University Göttingen, Göttingen Center for Molecular Biosciences (GZMB) and University Medical Center Göttingen (UMG), Institute of Molecular Oncology, Section for Cellular Oncology, D-37077 Göttingen, Germany; Department of Clinical Neurobiology, University Hospital Heidelberg and German Cancer Research Center (DKFZ), D-69120 Heidelberg, Germany; Genomics Core Facility, European Molecular Biology Laboratory (EMBL), Heidelberg, Germany; Institute of Human Genetics, Heidelberg University, D-69120 Heidelberg, Germany; MRC Laboratory of Molecular Biology, CB2 0QH Cambridge, UK; Department of Medical Biochemistry and Biophysics, Umeå University, 901 87 Umeå, Sweden; Nikon Imaging Center at the University of Heidelberg, Bioquant, D-69120 Heidelberg, Germany; Schaller Research Group, German Cancer Research Center (DKFZ) (DKFZ), D-69120 Heidelberg, Germany; Division of Molecular Thoracic Oncology, German Cancer Research Center (DKFZ) (DKFZ), D-69120 Heidelberg, Germany

**Author notes:** These authors contributed equally (A De Jaime-Soguero, J Hattemer). These authors contributed equally (H Bastians, SP Acebrón).

**Keywords:** Cell signalling, Chromosome segregation, DNA replication stress, DNA damage, Aneuploidy, BMP, WNT, lineage specification, pluripotent stem cells, neurogenesis

## Abstract

The development and homeostasis of organisms rely on the correct replication, maintenance and segregation of their genetic blueprints. How these intracellular processes are monitored across generations of different human cellular lineages, and why the spatio-temporal distribution of mosaicism varies during development remain unknown. Here, we identify several lineage specification signals that regulate chromosome segregation fidelity in both human and mouse pluripotent stem cells. Through epistatic analyses, we find that that WNT, BMP and FGF form a signalling “rheostat” upstream of ATM that monitors replication fork velocity, origin firing and DNA damage during S-phase in pluripotency, which in turn controls spindle polymerisation dynamics and faithful chromosome segregation in the following mitosis. Cell signalling control of chromosome segregation fidelity declines together with ATM activity after pluripotency exit and specification into the three human germ layers, or further differentiation into meso– and endoderm lineages, but re-emerges during neuronal lineage specification. In particular, we reveal that a tug-of-war between FGF and WNT signalling in neural progenitor cells results in DNA damage and chromosome missegregation during cortical neurogenesis, which could provide a rationale for the high levels of mosaicism in the human brain. Our results highlight a moonlighting role of morphogens, patterning signals and growth factors in genome maintenance during pluripotency and lineage specification, which could have important implications for our understanding on how mutations and aneuploidy arise during human development and disease.

**One sentence summary:** Developmental signals link genome maintenance to cell fate

## Introduction

Organismal viability requires faithful duplication and transmission of the genetic material across different cellular lineages. As such, intrinsic cellular mechanisms tightly monitor DNA replication and repair during S-phase, as well as chromosome segregation during mitosis ^1–4^. Despite these conserved cellular checkpoints, the rate and distribution of *de novo* genomic variations are neither low nor homogenous across all developmental lineages and cell types ^5–8^. In particular, the majority of human preimplantation embryos, and up to 30% of neural progenitors during mammalian brain development, present numerical chromosomal aberrations ^5, 9–12^, while other embryonic lineages display low levels of mosaicism. The existence of these developmental bottlenecks suggests that cell fate-dependent mechanisms might differentially affect chromosome segregation fidelity during human development.

Cell lineage specification is controlled by morphogens, patterning signals and growth factors that form distinct gradients to shape embryos, direct tissue patterning, and regulate cell fate ^13–17^. Critically, chromosomal instability (CIN) risks weakening the robustness of these developmental and cellular signalling programmes ^18–21^. Here, we investigate whether morphogens, patterning signals and growth factors can in turn regulate chromosomal stability during human lineage specification (Figure 1A).

**Figure 1:**
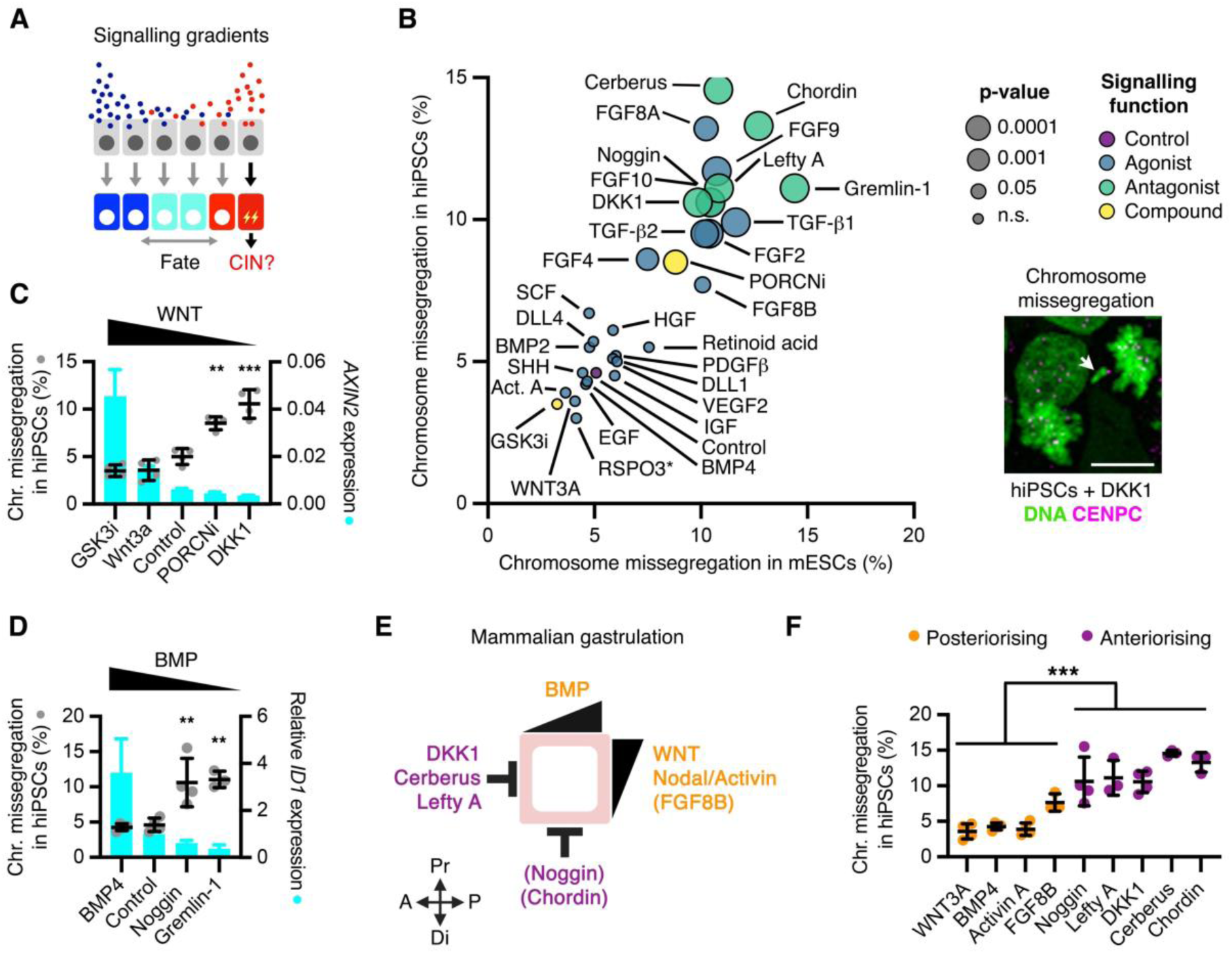
Embryo patterning signals regulate chromosome segregation fidelity in PSCs. **A**, Schematic of signalling-driven patterning and the hypothesised role in chromosomal instability (CIN). **B**, Chromosome segregation analyses in mESCs and hiPSCs upon treatment for 16 hours with different pathway-specific activating signals (agonists), inhibiting signals (antagonists), or small molecule compounds. Data are mean and *P-*value of 3 independent experiments with >100 anaphases per condition in each experiment. An example of a DKK1-treated hiPSC in anaphase is shown. DAPI stains the DNA and CENPC marks kinetochores. **C,D**, Chromosome missegregation from **(B)** and qRT-PCR analyses of the WNT target gene *AXIN2* **(C)** or the BMP target gene *ID1* **(D)** in hiPSCs upon treatment for 16 hours with the indicated compounds and proteins. **E**, Schematic of the signalling axes driving gastrulation in mammalian embryos. Note that Chordin and Noggin do not establish the anterior visceral endoderm in mammals, but are required for subsequent anterior patterning ^30^ (See also table 1). **F**, Chromosome segregation analyses in hiPSCs from **(B)** upon treatment with signals promoting anteriorisation or posteriorisation during mammalian gastrulation. *P*-values from t-test between all independent experiments from the indicated groups, **P < 0.01, ***P < 0.001.

To gain access and map signalling decisions across bifurcating fate choices from pluripotency to human lineage specification, we turn into primed pluripotent stem cells, which represent *bona fide* model of the epiblast before gastrulation. We show that developmental signals control chromosome segregation fidelity during pluripotency and neurogenesis, but not in other embryonic lineages, by modulating the cellular response to replicative stress. We identify that WNT and BMP signalling are at the helm of this regulatory cascade by protecting cells from different sources of DNA replication stress, including other signalling cascades and the perturbation of core DNA replication components. Given the spatio-temporal patterns of activation and co-occurrence of the examined signalling cascades during mammalian development, our findings could provide a rationale to understand why mosaicism is prevalent in early embryogenesis and the developing brain.

## Results

### Identification of signals controlling chromosome segregation in pluripotent stem cells

We have previously identified that WNT ligands promote chromosome segregation fidelity in somatic and cancer cells ^22–24^, although the underlying mechanisms and biological relevance remained uncharacterised. We hypothesised that WNT and possibly other developmental signalling pathways could exert a control of ploidy in targeted embryonic lineages (Figure 1A), thereby explaining spatio-temporal distribution of chromosomal mosaicism during development. To examine the roles of cell signalling on chromosome segregation fidelity during early lineage specification, we treated primed human induced pluripotent stem cells (hiPSCs) with morphogens, growth factors, and chemical inhibitors targeting the main developmental signalling pathways that regulate stem cell maintenance, lineage specification and embryonic patterning (Table 1). These factors and compounds were used at physiologically relevant conditions, i.e., by inhibiting endogenous signals or activating pathways at commonly used concentrations during cell lineage specification studies (Table 1). Beyond hiPSCs, we also analysed mouse embryonic stem cells (mESCs) to determine the conservation of the effects between mammals and across naive and primed pluripotency. We treated hiPSCs and mESCs with signalling modulators for 16 hours, and analysed chromosome missegregation rates by immunofluorescence in three independent experiments for a total of ∼18,000 anaphases (Figure 1B). Inhibition of autocrine WNT signalling by its physiological antagonist DKK1, inhibition of endogenous Bone Morphogenetic Protein (BMP) signalling using Noggin, Chordin or Gremlin 1, or inhibition of WNT, BMP and NODAL by Cerberus, increased >2-fold the chromosome missegregation rate in both mESCs and hiPSCs (Figure 1B). Furthermore, activation of Transforming Growth Factor (TGF) by TGF-β1/2, inhibition of Nodal signalling by LEFTY-A, as well as Fibroblast Growth Factor (FGF) signalling activation by various ligands (FGF2/4/8A/8B/9), also promoted chromosome segregation errors in mESCs and hiPSCs (Figure 1B). For factors and compounds regulating the same pathway, signalling activity levels correlated with their effects on chromosome segregation fidelity (Figure 1C and 1D), suggesting that signalling gradients could differentially impact chromosomal stability. Human PSCs represent a great model to study gastrulation *in vitro* ^25^. Intriguingly, we observed that signals promoting anteriorisation (DKK1, Noggin, LEFTY-A, Cerberus, Chordin) during mammalian gastrulation were strongly associated with higher chromosome missegregation rate in pluripotent stem cells, compared with signals promoting posteriorisation (WNT3A, BMP4, Activin A/Nodal, FGF8B) (Figure 1F, S1A-C, Table 1).

We decided to focus our studies on WNT, BMP, and FGF signalling (Figure 2A), which are critical drivers of lineage specification and embryonic patterning ^17, 25–33^. In addition to mESCs and hiPSCs (Figure 1B), we found that inhibition of endogenous WNT and BMP signalling using DKK1 and Noggin, respectively, as well as FGF signalling activation by FGF2, also induced chromosome missegregation in primed human embryonic stem cells (hESCs) (Figure S2A and S2B). To confirm the specificity of the investigated signalling cascades, we performed rescue experiments by targeting their downstream effectors. Inhibition of GSK3 using CHIR99021 (GSK3 inhibitor, GSK3i), which activates the canonical WNT pathway downstream of the receptor complex ^34^ (Figure 2A), rescued DKK1-induced chromosome missegregation in hiPSCs (Figure 2B). Furthermore, transient inhibition of the FGF receptor with a specific small compound (CAS 192705-79-6, FGFRi) (Figure 2A), or co-treatment with BMP4, rescued chromosome missegregation in hiPSCs induced by FGF2 and Noggin, respectively (Figure 2B). Treatment of hiPSCs with DKK1, FGF2, or Noggin neither impacted cell cycle progression nor induced differentiation priming after 16 hours (Figure S2C and S2D). Furthermore, triggering exit of pluripotency by placing cells in E6 or RPMI media for 16 hours did not result in increased chromosome missegregation (Figure S2E), indicating that signalling roles in chromosome segregation in hiPSCs are not due to cell differentiation or proliferation.

**Figure 2:**
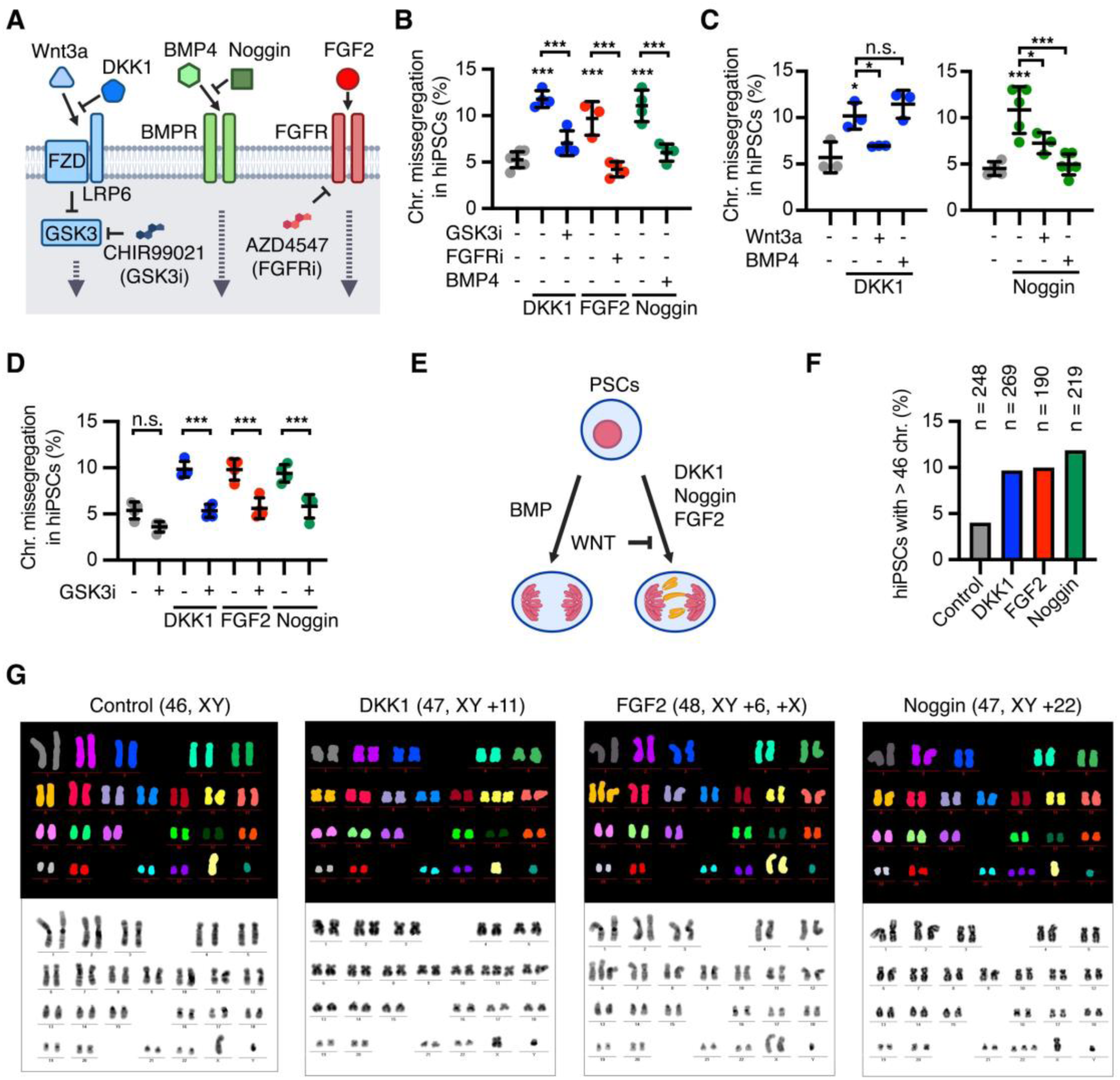
Embryo patterning signals regulate chromosome segregation fidelity in PSCs. **A**, Simplified scheme of the WNT, BMP and FGF signalling pathways highlighting the proteins and modulators used in this work. **B-D**, Chromosome segregation analyses in hiPSCs treated as indicated for 16 hours. Data are mean ± s.d. of n= 3-5 experiments with >100 mitotic cells per condition. *P*-values from one-way ANOVA analyses with Tukey correction are indicated as *P < 0.05, **P < 0.01, ***P < 0.001, or n.s., not significant. **E**, Schematic of the epistatic interactions between WNT, FGF and BMP signalling in chromosome segregation fidelity. **F**, Analyses of the chromosome gains in >150 Giemsa staining and 50 M-FISH analyses per condition are shown. **G**, M-FISH examples of hiPSCs treated as indicated.

Morphogenetic signals often crosstalk epistatically to exert their functions in lineage specification ^13, 15, 31^. We found that activation of the WNT pathway by WNT3A or GSK3i was sufficient to rescue not only DKK1, but also FGF2 and Noggin effects in chromosome missegregation (Figure 2C and 2D). On the other hand, additional epistastic experiments revealed that BMP4 was not able to rescue inhibition of endogenous WNT signalling activity (Figure 2C). Taken together, these results indicate that WNT, BMP, and FGF signalling functionally interact upstream of GSK3 to regulate chromosome segregation fidelity during human pluripotency (Figure 2E).

Live cell imaging analyses of hiPSCs labelled with SiR-DNA confirmed that hiPSCs undergoing chromosome missegregation upon exposure to DKK1, FGF2, or Noggin completed their division and survived (Figure S2F and S2G). Accordingly, analyses of metaphase spreads of hiPSCs using multiplex FISH (M-FISH) and Giemsa staining revealed abnormal karyotypes after one cell cycle in the presence of DKK1, FGF2, or Noggin (Figure 2F, 2G, S2H and S2I). These results show that while exogenous activation of FGF signalling can induce aneuploidy, autocrine WNT and BMP signalling prevent chromosome instability in hiPSCs.

## WNT, BMP and FGF regulate DNA replication stress during S-phase

To gain insight into the molecular mechanisms underlying the effects of WNT, BMP, and FGF signalling, we performed single-cell RNA sequencing of hiPSCs (Figure 3A). Gene ontology analyses of differentially upregulated genes upon endogenous WNT and BMP signalling inhibition, or FGF signalling activation, showed the highest enrichment score for factors promoting H3K4 methylation (7.5-fold, *P* = 6.5×10^−07^), which is associated with the mitigation of replication stress ^35, 36^, as well as DNA unwinding factors involved in DNA replication (7.4-fold, *P* = 3×10^−06^) and ATP-dependent chromatin remodelling factors (6.8-fold, *P* = 3.8×10^−08^) (Figure 2A and Table 2), suggesting a role of these pathways during DNA replication in S-phase. Intriguingly, i) pluripotent stem cells deal with elevated replicative stress under normal self-renewing conditions ^37, 38^, and ii) replicative stress is a major driver of structural and numerical chromosomal instability 39,40.

**Figure 3:**
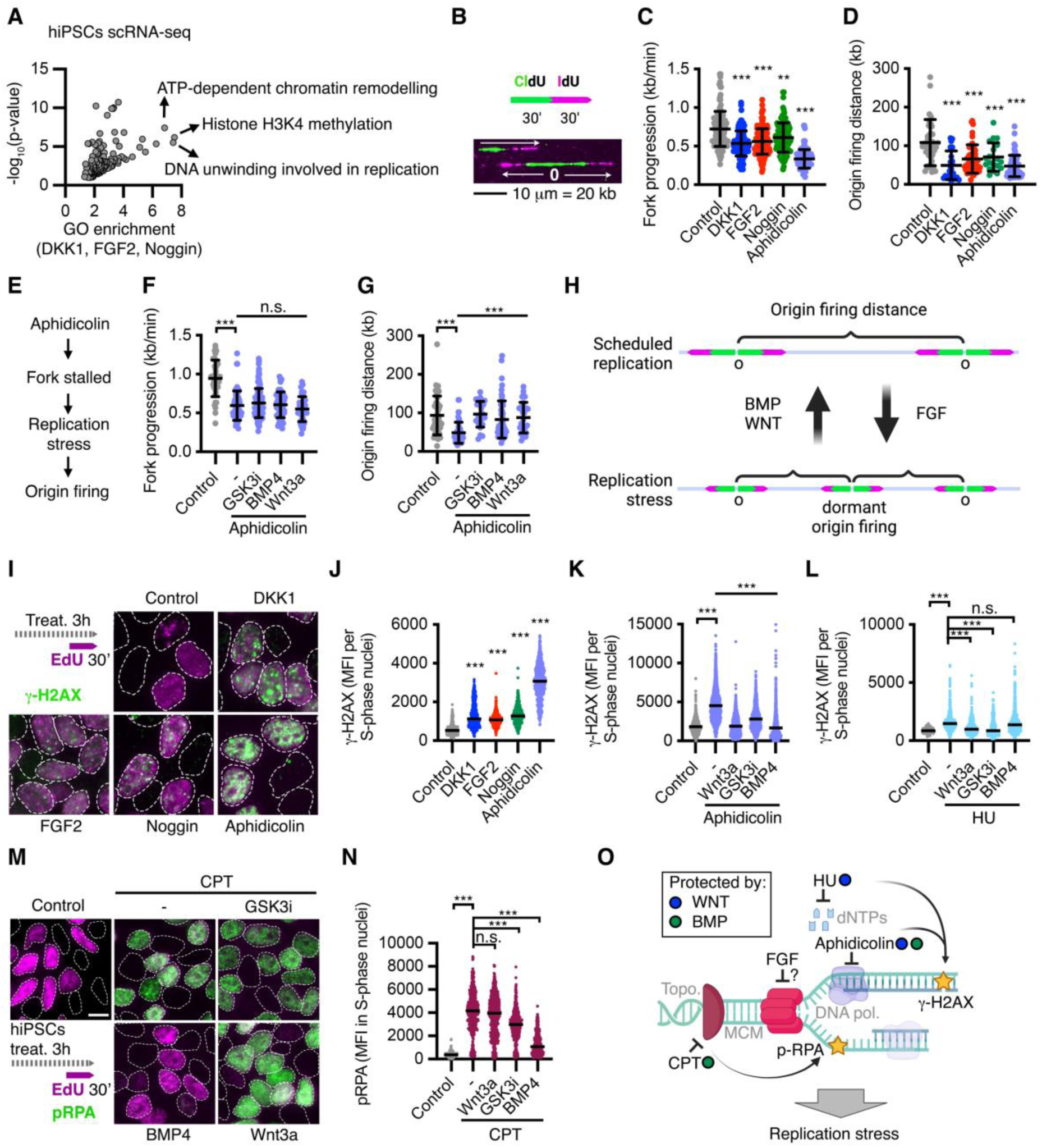
WNT, BMP and FGF converge into the regulation of DNA replication stress. **A**, Gene Ontology (GO) term enrichment analyses of common transcripts differentially upregulated by DKK1, FGF2, and Noggin in hiPSCs after 16 hours treatment. Expression analyses were performed by single cell RNA-sequencing of >75 cells per condition. **B-G**, DNA combing experiments in hiPSCs treated as indicated and labelled with consecutive pulses of CIdU and IdU as shown in **(B)**. Data are mean ± s.d of >100 single forks (**C,F**) or > 45 inter-origin distances (**D,G**). In **(B)** examples of fork progression and origin (O) are shown. In **(E)** a scheme of the replication stress cascade targeted in **(F,G)** is shown. **H**, Proposed roles of WNT, BMP and FGF signalling in DNA replication. **I-N**, Accumulation of γ-H2AX foci **(I-L)** and phospho-RPA **(M,N)** during S-phase in hiPSCs treated for 3h as indicated. HU, hydroxyurea; CPT, camptothecin. Data are median fluorescence intensity (MFI) at EdU^+^ nuclei of representative experiments performed at least 3 independent times with > 150 cells per condition in each experiment. **O**, Schematic summarising the signalling functions of WNT, BMP and FGF upon DNA replication stress. *P*-values from one-way ANOVA analyses with Tukey correction are indicated as *P < 0.05, **P < 0.01, ***P < 0.001, or n.s., not significant.

To examine the possibility that WNT, BMP, and FGF signalling function in DNA replication, we performed single molecule analyses of replication dynamics in hiPSCs by DNA combing (Figure 3B). Inhibition of WNT and BMP signalling, as well as activation of FGF signalling, reduced replication fork velocity in hiPSCs between 16-25% (Figure 3C). Slowed replication rates are associated with DNA replication stress response, which includes the firing of dormant origins as a compensatory mechanism to facilitate completion of DNA replication in eukaryotes ^41^. Consistently, we found that DKK1, FGF2, and Noggin decreased the distance between origin firings from ∼100 kb to 70-49 kb (Figure 3D), similar to treatment with low dose of the DNA polymerase inhibitor aphidicolin (Figure 3D), which stalls replication forks. Conversely, co-treatment with WNT3A, GSK3i, or BMP4 rescued origin firing induced by aphidicolin without affecting replication fork velocity (Figure 3E, 3F and 3G), which indicates a role of WNT and BMP signalling in the mitigation of DNA replication stress downstream of stalled forks (Figure 3H). Unresolved DNA replication stress triggers subsequent errors, including double strand breaks (DSBs). Phosphorylation of histone H2AX (γ-H2AX) is an early ATM-dependent response to DSBs ^42, 43^, including after DNA replication stress in pluripotent stem cells ^44^. Inhibition of autocrine WNT and BMP signalling with DKK1 and Noggin, respectively, as well as FGF activation with FGF2 for 3 hours induced the formation of γ-H2AX foci in hiPSCs specifically during S-phase (Figure 3I, 3J EdU^+^ cells), which is consistent with replication stress-associated DNA damage.

To get additional functional information on how WNT, BMP and FGF signalling impact DNA replication, we performed phospho-proteomics and epistatic experiments upon acute perturbation of these pathways. We treated hiPSCs for 3 hours with DKK1, Noggin and FGF2 in 4 independent experiments and analysed their phospho-proteome (Figure S3A). Among the most significantly regulated phospho-peptides across the treatments, we identified several factors associated with DNA replication and damage response (Figure S3B). Five phospho-peptides were differentially regulated in the three treatments (Figure S3C), including from the key DNA replication stress-associated proteins RIF1 ^40, 45, 46^, and BRCA1 at the CDK1-target serine 1191, which is triggered by DNA replication stress to stabilise it ^47, 48^ (Figure S3B-E). In addition to RIF1 and BRCA1, i) FGF activation differentially regulated factors directly controlling replisome progression (Figure S3B and S3D), including the downregulation of the phosphorylation of the replication initiation factor MCM2 at the CDC7-target serine 26 ^49^ (Figure S3B, S3D and S3E); ii) BMP signalling inhibition triggered phosphorylation of EEPD1 and the DNA chaperone HMGA2, which are associated with the stability of single stranded DNA (ssDNA) at stalled forks ^50, 51^, and affected several factors functioning at the DNA damage response (Figure S3B and S3D); and iii) DKK1 differentially regulated the phosphorylation of XRNA2 and SF3B1, which function at the fork in resolving R-loops during DNA replication stress ^52, 53^, as well as additional factors involved in the DNA damage response, including TP53BP1 (Figure S3B and S3D). Taken together, these results suggest that FGF signalling may function during DNA replication initiation and/or progression, while BMP and WNT might have distinct roles in the modulation of DNA replication stress and DNA damage response, which we decided to further explore through further epistatic analyses.

In agreement with our DNA combing experiments (Figure 3C-3G), WNT and BMP activation – but not FGF signalling inhibition by FGFRi (Figure S4A) – impaired the formation of γ-H2AX foci upon partial inhibition of the DNA polymerase using aphidicolin (Figure 3K). Intriguingly, additional rescue experiments indicated that WNT, but not BMP signalling, protect hiPSCs in S-phase from DSBs upon acute dNTP depletion using hydroxyurea (HU) (Figure 3L). Topoisomerase I inhibition by camptothecin (CPT) triggers recruitment and phosphorylation of the single strand DNA break proteins RPA1-3 (p-RPA) at the stalled forks ^54^, which was rescued by BMP signalling activation, but not by WNT signalling (Figure 3M and 3N). These results, together with our phospho-proteomics (Figure S3) and previous epistatic analyses (Figure 2C, 2D) support a role of FGF signalling interfering with DNA replication initiation or progression, a function of BMP signalling in protecting ssDNA at the stalled replication forks, and a downstream role of WNT signalling in successfully resolving DNA replication stress (Figure 3O). Future molecular studies are required to disentangle the specific targets and functions of either pathway across the different steps associated with replicative stress.

## WNT, BMP and FGF signalling regulate chromosome segregation in mitosis through the modulation of DNA replication stress in the previous S-phase

In addition to DNA damage, recent evidence demonstrated that replicative stress can induce chromosome missegregation in somatic cells through CDC7-, and CDK1-dependent elevated microtubule dynamics in the subsequent mitosis (Figure 4A) ^40^. Interestingly, loss of WNT signalling also upregulates polymerisation of microtubule plus-ends in somatic cells ^23^. Live cell imaging analyses of EB3-eGFP in hiPSCs revealed that DKK1, Noggin, and FGF2 increased microtubule polymerisation dynamics during mitosis (Figure 4A and 4B). To determine whether cell signalling effects in the mitotic spindle were caused by their roles in DNA replication, we co-treated cells with nucleosides (dNs), which alleviate DNA replication stress and prevent DNA replication-associated damage ^55, 56^. Treatment of hiPSCs with nucleosides indeed rescued the upregulation of microtubule polymerisation by DKK1, Noggin, and FGF2 (Figure 4B), supporting a mechanistic link between WNT, BMP, and FGF roles in S– and M-phase.

**Figure 4:**
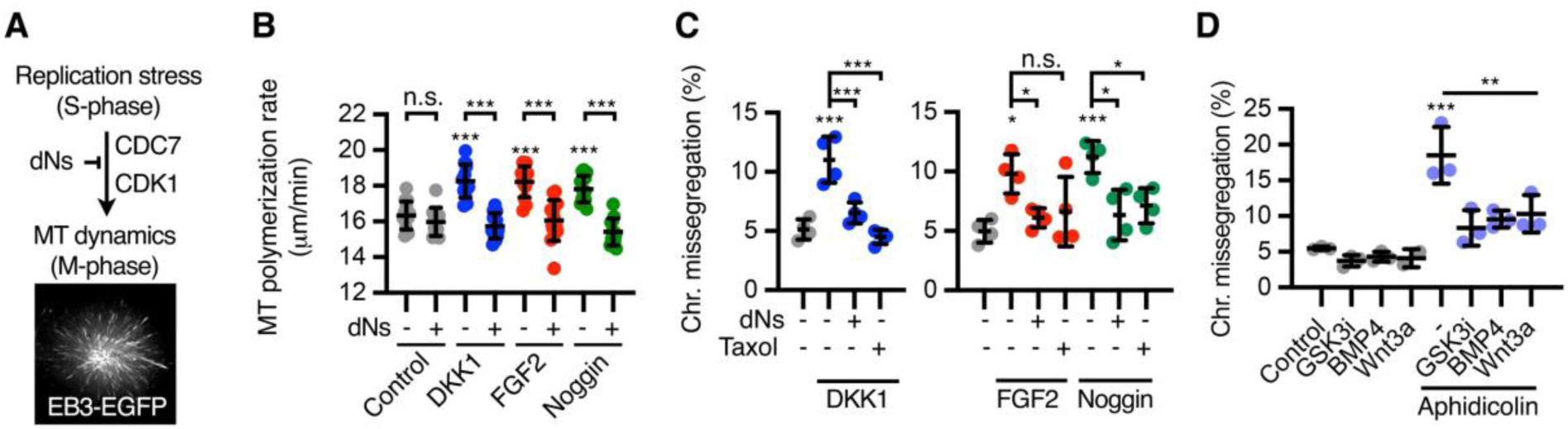
WNT, BMP and FGF control chromosome segregation through their roles in replication. **A,B**, Mitotic microtubule plus-end assembly rates measured by EB3-GFP tracking during prometaphase in hiPSCs. Cells were treated as indicated and arrested in mitosis using dimethylenastrone. Data are mean ± s.d. of average growth rates of 20 microtubules/cell. **C,D**, Chromosome segregation analyses in hiPSCs treated as indicated for 16 hours. Data are mean ± s.d. of n = 3-4 experiments with >100 mitotic cells per condition. *P*-values from one-way ANOVA analyses with Tukey correction are indicated as *P < 0.05, **P < 0.01, ***P < 0.001, or n.s., not significant for single cells of a representative experiment **(B)**, or independent experiments **(C,D)**.

Critically, attenuation of replicative stress with nucleosides, as well as stabilisation of microtubule dynamics using a low dose of taxol, rescued chromosome missegregation induced by DKK1, FGF2, or Noggin in hiPSCs (Figure 4C). Furthermore, and consistent with our replication stress analyses (Figure 3G and 3K), exogenous activation of WNT or BMP signalling rescued aphidicolin-induced chromosome missegregation in hiPSCs (Figure 4D). Taken together, our data indicates i) that FGF signalling induces replicative stress-dependent chromosome missegregation in pluripotent stem cells, and ii) that autocrine WNT and BMP signalling promote genome stability and chromosome segregation fidelity in pluripotent stem cells by preventing excessive origin firing downstream of stalled forks during S-phase (Figure 3O). In the case of WNT signalling, our findings further highlight a complex programme driven by this pathway to promote faithful G1, S– and M-phase progression ^23, 24, 57–59^.

## Limited function of morphogens in chromosomal stability after germ layer specification

During human embryonic development, WNT3A and NODAL direct epiblast cells into a transient posterior structure named primitive streak prior commitment to the mesoderm and the definitive endoderm, a process that can be mimicked *in vitro* using WNT3A and Activin A (Figure S5A and S5B) ^27, 29^. In contrast to pluripotent stem cells, inhibition of WNT signalling did not result in chromosome missegregation in human primitive streak-like cells (Figure 4A and Figure S5A-C), and had a limited effect in the definitive endoderm-like cells (Figure 5A, S5A, S5D and S5E). Upon FGF induction, WNT and BMP signalling display antagonistic functions in the primitive streak differentiation into paraxial and lateral mesoderm (Figure S5A and S5F) ^27^. Perturbation of WNT, BMP or FGF signalling during the last 16 hours of the specification of these lineages did not impact chromosome segregation fidelity (Figure 5A and S5G). To get insight into signalling roles in the ectoderm lineage, we turned into neuroectoderm specification, which is the first step in the development of the nervous system ^15^. We performed Noggin-directed differentiation of hiPSCs into neuroectoderm-like cells (Figure S5A and S5H) ^28, 60^. Neuroectoderm cells displayed slightly higher levels of chromosome missegregation (6%) compared to primitive streak, endoderm, or mesoderm (∼3%) (Figure S5G and S5I). Although treatment with BMP4 slightly reduced chromosome missegregation in neuroectoderm-like cells, Noggin withdrawal or DKK1 treatment had no effect (Figure 5A and S5I). We also generated neural crest-like cells and early neural progenitors (NPCs) by a pairwise decision through activation or inhibition of WNT activity, respectively (Figure S5A and S5J) ^61^. Early neural progenitors displayed higher basal chromosome missegregation than neural crest-like cells (Figure S5K), but perturbation of WNT signalling had no significant impact in chromosome segregation in either lineage (Figure 5A and S5K). These results indicate that WNT, BMP, and FGF roles in chromosome segregation fidelity are largely impaired after specification into the three human germ layers, regardless of their critical signalling functions in cell fate determination in those lineages ^27, 28, 60^ (Figure S5A). In contrast to FGF2, DKK1, or Noggin – and despite the presence of WNT3A (Figure S5A) –, aphidicolin treatment induced DNA damage and chromosome missegregation in primitive streak, endoderm, and mesoderm-like cells (Figure 5A, S5L and S5M). This indicates that specified cells display a functional DNA replication stress response but it is uncoupled of extracellular signalling activity. Aphidicolin did not induce high levels of γ-H2AX foci or chromosome missegregation in neuroectoderm, early neural progenitor cells, or neural crest cells (Figure 5A, S5L and S5M), possibly due to additional mechanisms protecting from DNA replication stress in these cells.

**Figure 5:**
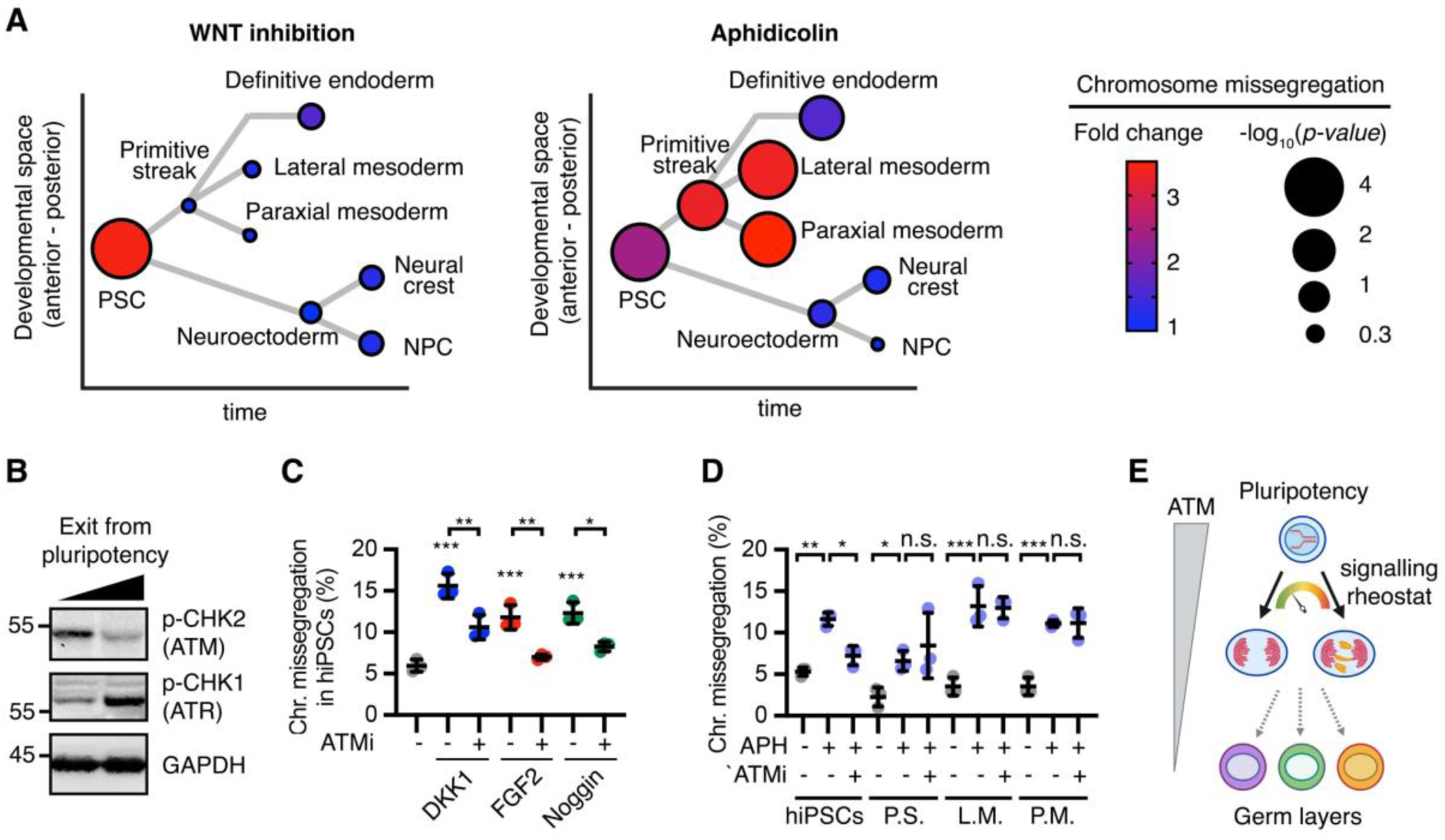
WNT, BMP and FGF do not impact chromosomal stability after human early lineage specification. **A**, Summary of chromosome segregation analyses across different embryonic lineages obtained upon differentiation of hiPSCs as further described in Supplementary Figure 5a. Data show fold changes in chromosome missegregation rates upon DKK1 treatment (WNT inhibition) or aphidicolin and *P*-values of 3-7 biological replicates per experiment shown in Supplementary Figure 5. Data are displayed in a lineage tree depicting relative developmental position in time of each lineage. NPC, neural progenitor cell. **B**, Western blot analyses of hiPSCs undergoing differentiation towards primitive streak. The molecular weight markers are indicated in kDa. **C,D**, Chromosome segregation analyses in hiPSCs, primitive streak-like (P.S.), lateral mesoderm (L.M.) and paraxial mesoderm (P.M.) –like cells treated as indicated for 16 hours. APH, 50 nM aphidicolin. ATMi, 3 μM AZD0156. Data are mean ± s.d. of 3 biological replicates with >100 mitotic cells per condition. *P*-values from one-way ANOVA analyses with Tukey correction are indicated as *P < 0.05, **P < 0.01, ***P < 0.001, or n.s., not significant. **E**, Schematic of the functional interaction between signalling roles in chromosome segregation and cell fate.

ATM signalling regulates the DNA replication stress response in pluripotent stem cells ^38^, in addition to the most common ATR response. We observed that exit from pluripotency towards primitive streak in human cells is accompanied by reduced phosphorylation of the ATM target CHK2, and increased phosphorylation of the ATR target CHK1 (Figure 5B). We hypothesised that a switch between ATM and ATR signalling could mediate the cell fate-dependent roles of morphogens in chromosomal stability. Indeed, inhibition of ATM signalling using AZD0156 (ATMi) rescued chromosome missegregation induced by DKK1, Noggin, and FGF2, as well as aphidicolin, in hiPSCs (Figure 5C and 5D). However, ATMi did not rescue chromosome missegregation induced by aphidicolin in human primitive streak, lateral mesoderm or paraxial mesoderm (Figure 5D). Taken together, these results indicate that the effects of WNT, BMP and FGF signalling on chromosome segregation fidelity decline after lineage specification in the three human germ layers following the withdrawal of ATM signalling as first responder to DNA replication stress (Figure 5E).

## A tug-of-war between WNT and FGF controls chromosomal stability during neurogenesis

Beyond peri-implantation and gastrulating embryos, only the neocortex displays high levels of genomic and chromosomal mosaicism later during embryonic development and adulthood ^5, 12, 62–65^.

To study the roles of cell signalling in chromosome segregation fidelity across more differentiated lineages, we generated cardiomyocyte-like (mesoderm), hepatocyte-like (endoderm), and neuronal progenitor cells (ectoderm) from hiPSCs (Figure S6A-J). Despite their response to chemically-induced replicative stress, neither cardiomyocyte nor hepatocyte-like cells at different stages of differentiation showed increased chromosome missegregation upon WNT inhibition (Figure S6C and S6G), in sharp contrast with neural progenitors, as further explained below.

To model human cortical neurogenesis *in vitro*, we differentiated hiPSCs into neural progenitors (hiNPCs) during a 16-day course (Figure 6A, 6B and S6H) ^66, 67^. Neural progenitor proliferation and neurogenesis are promoted by FGF and WNT signalling, among other pathways ^68–72^. *In vitro* generated hiNPCs expressed primarily *WNT8B*, in addition to *WNT1/3A/10B* (Figure S6I). Inhibition of autocrine WNT signalling activity by DKK1, as well as activation of FGF signalling with FGF2, strongly increased chromosome missegregation in hiNPCs during neurogenesis (day 16), but had no effect during hiNPC expansion (day 10) (Figure 6C, 6D and S6J). DNA combing and γ-H2AX foci analyses confirmed that both FGF activation and WNT inhibition promoted DNA replication stress and DNA damage in differentiating hiNPCs (Figure 6E-G). Alleviation of replicative stress in hiNPCs using nucleosides rescued chromosome missegregation induced by DKK1 and FGF2 (Figure 6H, I). Furthermore, exogenous activation of the WNT pathway by GSK3i or WNT3A during *in vitro* neurogenesis rescued FGF2– and aphidicolin-induced chromosome missegregation (Figure 6I, J).

**Figure 6:**
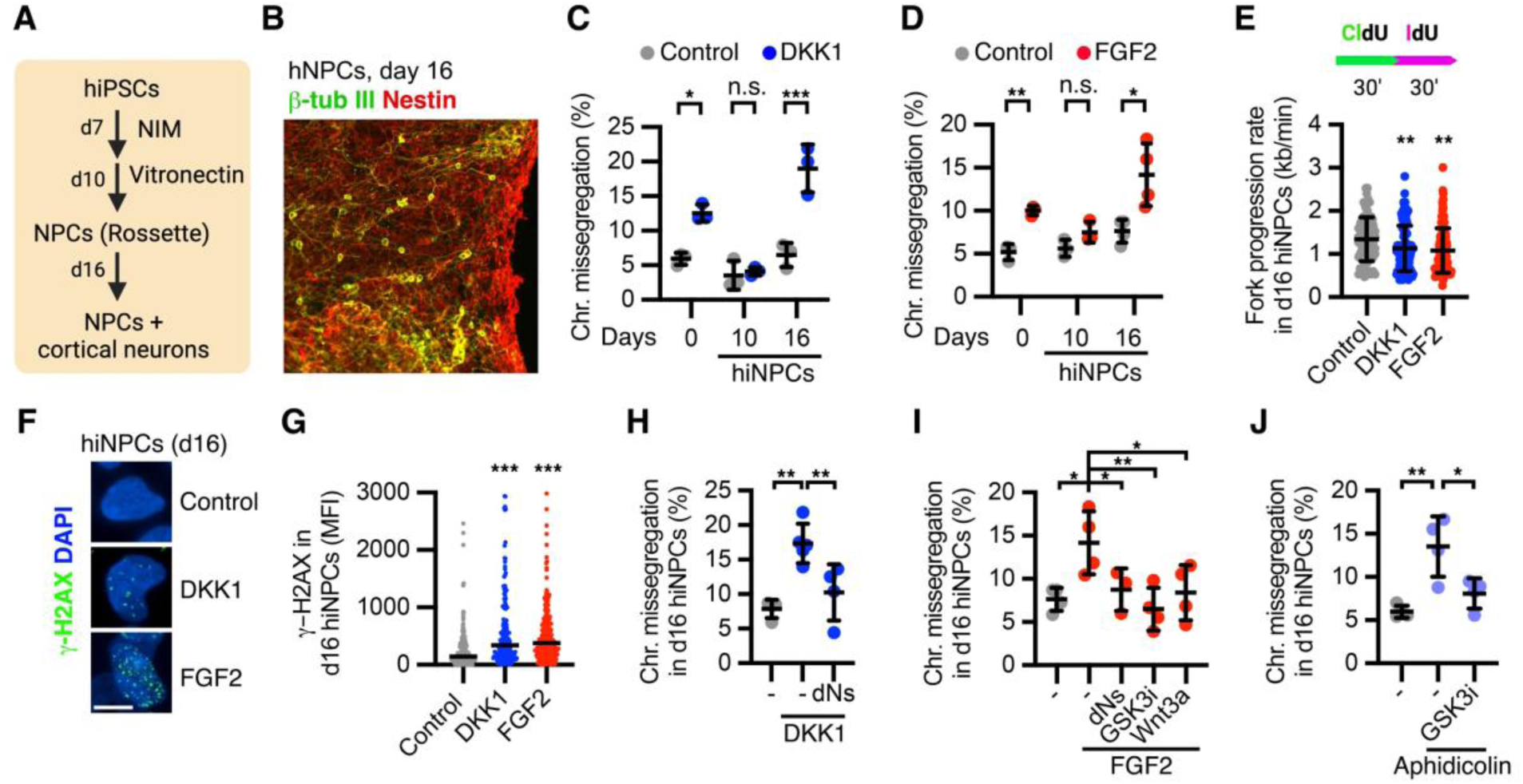
WNT and FGF display antagonistic roles in genome stability during human *in vitro* neurogenesis. **A,B**, *In vitro* specification of hiPSCs into cortical neural progenitors and neurons. **C,D**, Chromosome segregation analyses in hiPSCs, expanding hiNPCs (10 days) and differentiating hiNPCs (16 days) as indicated in (**A**), and treated with DKK1 or FGF2 16 hours before harvesting. **E**, DNA combing experiments in differentiating hiNPCs treated as indicated for 3h, and labelled with consecutive pulses of CIdU and IdU. Data are mean ± s.d. of > 80 single forks. **F,G**, Accumulation of γ-H2AX foci in differentiating hiNPCs treated as indicated. Data are median of the fluorescence intensity (MFI) in > 200 nuclei. **H-J**, Chromosome segregation analyses in differentiating hiNPCs treated as indicated for 16 hours. Data are mean ± s.d. of 4 biological replicates with 50-100 mitotic cells per condition.

Next, we used mouse embryonic brains and primary neural progenitors (NPCs) to determine whether our findings translate to *in vivo* and *ex vivo* conditions. In the developing mouse neocortex, NPCs expand at the ventricular zone before E13.5, followed by direct and indirect neurogenesis during E13.5-E15.5 ^68, 73^. Expression of WNT ligands, as well as phosphorylation of the WNT receptor LRP6, remained mostly constant in the developing ventricular zone during E12.5-E16.5 (Figure 7A, 7B, S7A and S7 B). *Fgf2* and *Fgfr1* are expressed in the ventricular zone of rodents before the onset of neurogenesis ^74^. Accordingly, we found that FGFR1 receptor activation peaked in the apical progenitors during E14.5 (Figure 7A, C). We hypothesised that a differential ratio between FGF and WNT activity during neurogenesis might contribute to the observed high levels of mosaicism in the developing brain ^5, 12, 62–65^.

**Figure 7:**
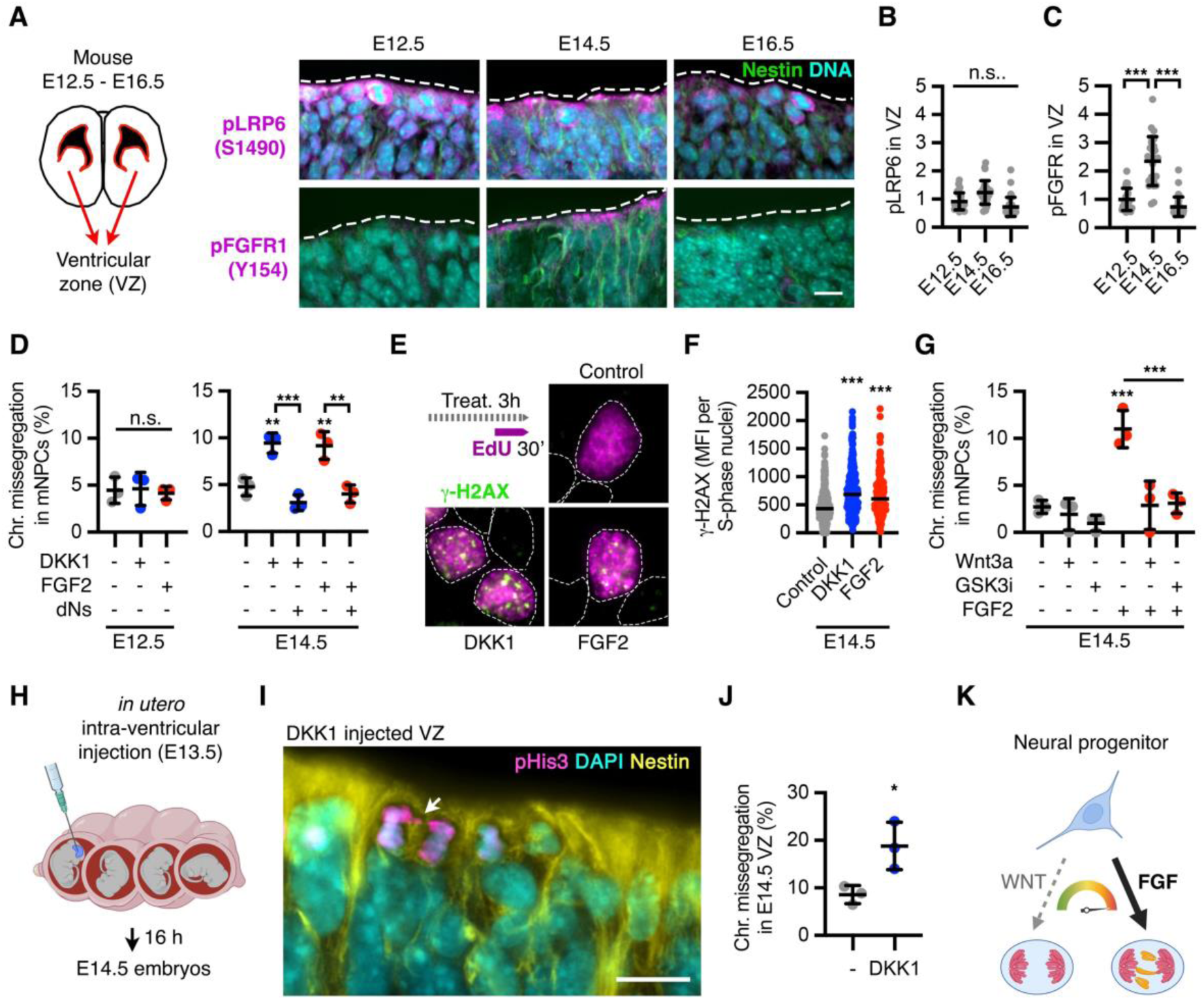
A tug-of-war between WNT and FGF regulates chromosome segregation fidelity in mouse neural progenitors. **A-C**, Immunofluorescence analyses of the activated WNT (LRP6) and FGF (FGFR1) receptors in the ventricular zone of E12.5-16.5 embryonic mouse brains. Quantification of phospho-LRP6 (S1490) **(B)** and phospho-FGFR1 (Y154) **(C)** in NPCs from the ventricular zone of mouse embryos is shown for three developmental stages of cortical neurogenesis. Data are mean ± s.d. of the relative fluorescence intensity normalised to Nestin in apical neural progenitors of > 5 brain cryosections. **D**, Chromosome segregation analyses in *ex vivo* cultured NPCs from E12.5 and E14.5 mouse embryos, treated as indicated for 16 hours. Data are chromosome missegregation rates from 3 independent experiments with >50 anaphases per condition. **E,F**, Accumulation of γ-H2AX foci in S-phase of E14.5-derived NPCs treated for 3h as indicated. Data are median of the fluorescence intensity (MFI) in > 200 EdU+ nuclei. **G**, Chromosome segregation analyses in *ex vivo* cultured NPCs from E14.5 mouse embryos, treated as indicated for 16 hours. Data are chromosome missegregation rates from 3 independent experiments with 50-100 anaphases per condition. **H**, Schematics of *in utero* ventricular injections of PBS (Control) or recombinant DKK1 in E13.5 mouse embryos, later sacrificed at stage E14.5. **I,J**, Chromosome segregation analyses in the ventricular zone of PBS (Control) or DKK1 injected mouse E14.5 embryos. Data are mean ± s.d. of 3 injected embryos per condition (>10 cryosections per embryo). **K**, A tug-of-war between WNT and FGF controls chromosome segregation fidelity in NPCs, and might underlie the high levels of chromosome missegregation occurring during neurogenesis. *P*-values from one-way ANOVA analyses with Tukey correction are indicated as *P < 0.05, **P < 0.01, ***P < 0.001, or n.s., not significant.

To study the functional interaction between WNT and FGF signalling, we isolated NPCs from E12.5 and E14.5 mouse embryonic cortices, which mainly expressed *Wnt7a/b* (Figure S7A and S7B). Inhibition of endogenous WNT signalling by DKK1 or activation of FGF signalling using FGF2 induced chromosome missegregation in E14.5-derived NPCs, but not in E12.5-derived NPCs (Figure 7D, S7B and S7C). These results echoed our finding in expanding vs. differentiating human iNPCs (Figure 6C and 6D), further supporting a selective conserved role of WNT and FGF signalling in controlling chromosome segregation fidelity during cortical neurogenesis. WNT and FGF antagonising roles in chromosome segregation fidelity were also tracked back to S-phase in mouse primary NPCs: first, alleviation of DNA replication stress using nucleosides rescued chromosome segregation defects induced by DKK1 and FGF2 (Figure 7D). Second, both FGF2 and DKK1 increased the number of γ-H2AX foci during S-phase (Figure 7E and 7F). As in the case of hiNPCs (Figure 6I), exogenous activation of WNT signalling with WNT3A or GSK3i rescued FGF2-induced chromosome missegregation in primary NPCs isolated from E14.5 mouse embryos (Figure 7G).

To characterise WNT signalling roles in chromosome segregation directly *in vivo*, we performed *in utero* intra-ventricular injections of recombinant DKK1 in E13.5 embryos and analysed their cortices at E14.5 (Figure 7H). We observed high levels of chromosome missegregation (9%) in the control embryos (Figure 7H-K, PBS injected), similar to previous estimates ^11, 12, 20, 62, 75^. Critically, *in vivo* inhibition of WNT signalling induced chromosome missegregation in 19% of dividing NPCs during embryonic neurogenesis (Figure 7J and 7J).

Finally, we enquired whether signalling-induced chromosome missegregation in NPCs impact their viability and fate. Tracking analyses of live cell imaging showed that chromosome missegregation in mouse E14.5-derived NPCs increased 2-fold the rate of both cell death (14% after a normal mitosis vs 30% after chromosome missegregation) and of additional segregation errors (5% after a normal mitosis vs 12% after chromosome missegregation) in the daughter cells, but surprisingly did not substantially impact the ratio of dividing/non-dividing daughter cells (Figure S7D-G). These results indicate that chromosome missegregation is largely tolerated in primary NPCs committed to neurogenesis and could contribute to chromosomal mosaicism in the brain.

In summary, our findings show that a tug-of-war between the WNT and FGF pathways controls chromosome segregation fidelity during mouse and human neurogenesis (Figure 7K).

## Discussion

Morphogens, patterning signals and growth factors direct cells to build and maintain tissues, organs and organisms by interpreting genotypes into gene-regulatory networks. In this study, we show that developmental signals not only *read* genetic blueprints, but also play a moonlighting role in their *maintenance*. We reveal a dichotomy between signals inducing anteriorisation and posteriorisation during gastrulation in inducing or protecting pluripotent stem cells from chromosome missegregation, respectively. We identified that these patterning signals regulate DNA replication dynamics in S-phase and chromosome segregation fidelity in the following mitosis, and that WNT and BMP signalling sit at the helm of this regulatory cascade by protecting cells from different sources of replicative stress. Future studies are required to unravel the complexity of the replication mechanisms controlled by each signalling cascade. By performing *in vitro* lineage specification experiments with human pluripotent stem cells, we showed that cells rely on extracellular signals for faithful chromosome segregation during early lineage specification (before exit from pluripotency) and during neurogenesis, but not in specified human three germ layers or their subsequent differentiated lineages. Using mouse and human neural progenitors, we found that FGF to WNT signalling ratio determines chromosome missegregation rates during neurogenesis.

Predictions of this model are as follows: **1)** Morphogens and patterning signals should interlink DNA replication dynamics, DNA damage, and chromosomal stability with cell fate. Indeed, mutational and aneuploidy rates largely overlap during human development, peaking before gastrulation and during neurogenesis ^5, 9, 11^. Furthermore, DNA replication speed, ATM, γ-H2AX, and aneuploidy have recently emerged as key regulators of cell fate ^19, 76–78^, supporting a bi-directional relationship between them. Hence, it would be important to explore whether signalling roles in DNA replication provide an additional layer of control for lineage specification. **2)** Spatio-temporal signalling gradients driving the anterior-posterior axis in the gastrulating embryo should also generate a mosaicism gradient overlapping with the former. Human blastocysts do indeed show spatial chromosomal mosaicism ^79^, but it remains to be determined i) whether DKK1, Chordin, Cerberus, Noggin, and/or LEFTY-A induce higher levels of chromosomal instability in the anterior part of the embryo during gastrulation, ii) if NODAL, BMP4 and WNT3A ^29^ protect the organiser from replication stress and chromosome missegregation, and iii) whether these gradients trigger BMP4– and lineage-specific depletion of aneuploid cells ^7, 80^. **3)** Genomic alterations in the brain should be largely traced to neurogenesis, and not to e.g., neuroectoderm or expanding neural progenitors. This is the case ^5, 10, 11, 62^. Of note, genomic mosaicism has been proposed to foster neuronal specialisation ^81, 82^. It remains to be tested whether FGF-driven DNA replication stress and damage could render diversification in neuronal lineages at the cost of aneuploidy. In that respect, microglia are highly active during neurogenesis to remove aneuploid cells ^20^, and specific mitotic gene variants have evolved to partially reduce chromosome missegregation in the human developing brain ^83^. Given the technical and ethical challenges in testing these predictions in human embryos, the development of *in vitro* models of human gastrulation ^25, 80^ and brain development ^83–85^ opens now the opportunity to address these questions.

Our results, together with recent insights on SHH signalling ^86^, hypoxia ^87^, pathogens ^88^, and tissue architecture ^89^, highlight the importance of extracellular cues and cell fate for the correct replication, maintenance and segregation of chromosomes in mammalian cells.

## Methods

The detailed materials and methods can be found in the supplemental method section

## Data availability

Single cell sequencing datasets generated during this study will be available at ENA (*Number pending*)

## Ethics declarations

The authors declare no competing interests.

## Acknowledgements

We thank A. Smith, K.M. Noh, G. Pereira, M. Lancaster, and C. Niehrs for sharing reagents and cells. We thank the Nikon Imaging Center and the FACS facility at the University of Heidelberg for access to microscopes, cytometers, and for technical help. We thank the EMBL sequencing facility for technical support and sequencing analyses. This work was funded by the Deutsche Forschungsgemeinschaft (DFG, German Research Foundation) – SFB 1324 – Project number 331351713 (Project B03 to S.P.A. and H.B.). A.d.J.-S. held a Humboldt Research Fellowship for Postdoctoral Researchers. J.H. is recipient of a Studienstiftung PhD fellowship. J.A. and A.P. were supported by the Chica and Heinz Schaller Foundation. M.S. was supported by the Medical Research Council, as part of UK Research and Innovation (grant reference MRC, MC_UP_1201/24). V.S.R is supported by a Milstein fellowship.

## Author contributions

H.B. and S.P.A. designed the research; A.d.J.-S., J.H., A.B., and S.P.A. conceived, performed and analysed experiments. A.H., B.d.M., N.B., J.J.M.L., V.S.R., L.V., U.E, Y.B., B.D., S.A., M.T., Y.C.L., and A.J., performed and analysed experiments. V.B., M.S., A.J., A.P., R.S., and H.B. contributed new reagents/analytic tools and supervised aspects of the project. J.J.M.L and J.B., analysed the single cell sequencing data. S.P.A. supervised the project and wrote the paper, with input of the other authors.

## Supplementary Figures

**Figure S1:**
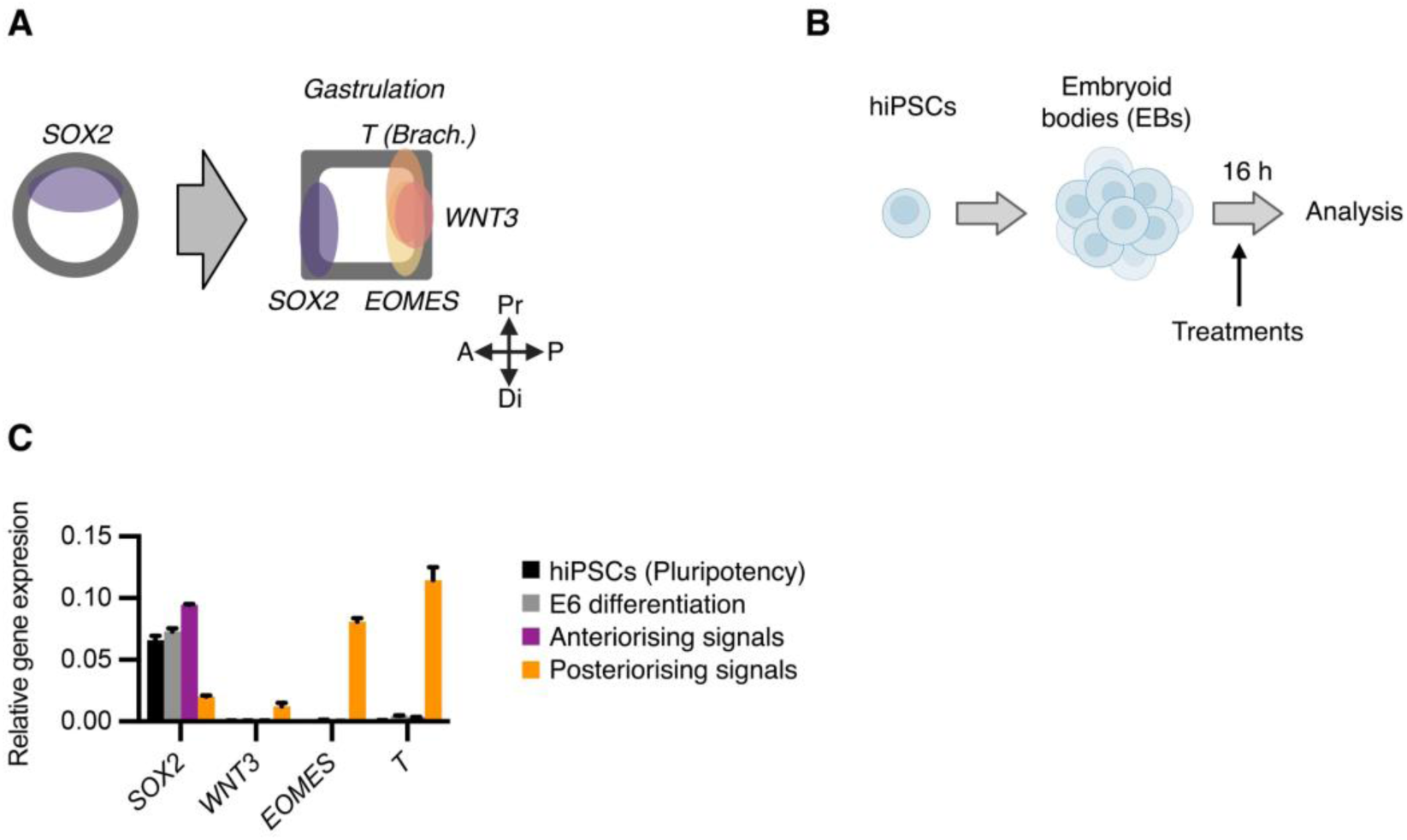
Morphogens modulate chromosome segregation fidelity in pluripotent stem cells. **A**, Model for the expression patterns of SOX2, EOMES, WNT3 and Brachyury (T) in mammals. **B**, Scheme of lineage specification experiment in embryoid bodies. **C**, Representative experiment of a qRT-PCR of hiPSCs cultured in E8 media (pluripotency) and hiPSC-derived embryoid bodies treated for 16 hours with E6 media (differentiation), E6 media supplemented with the anteriorising signals Noggin, Lefty A, DKK1, Chordin and Cerberus, or E6 media supplemented with the posteriorising signals Wnt3a, BMP4 and Activin A.

**Figure S2:**
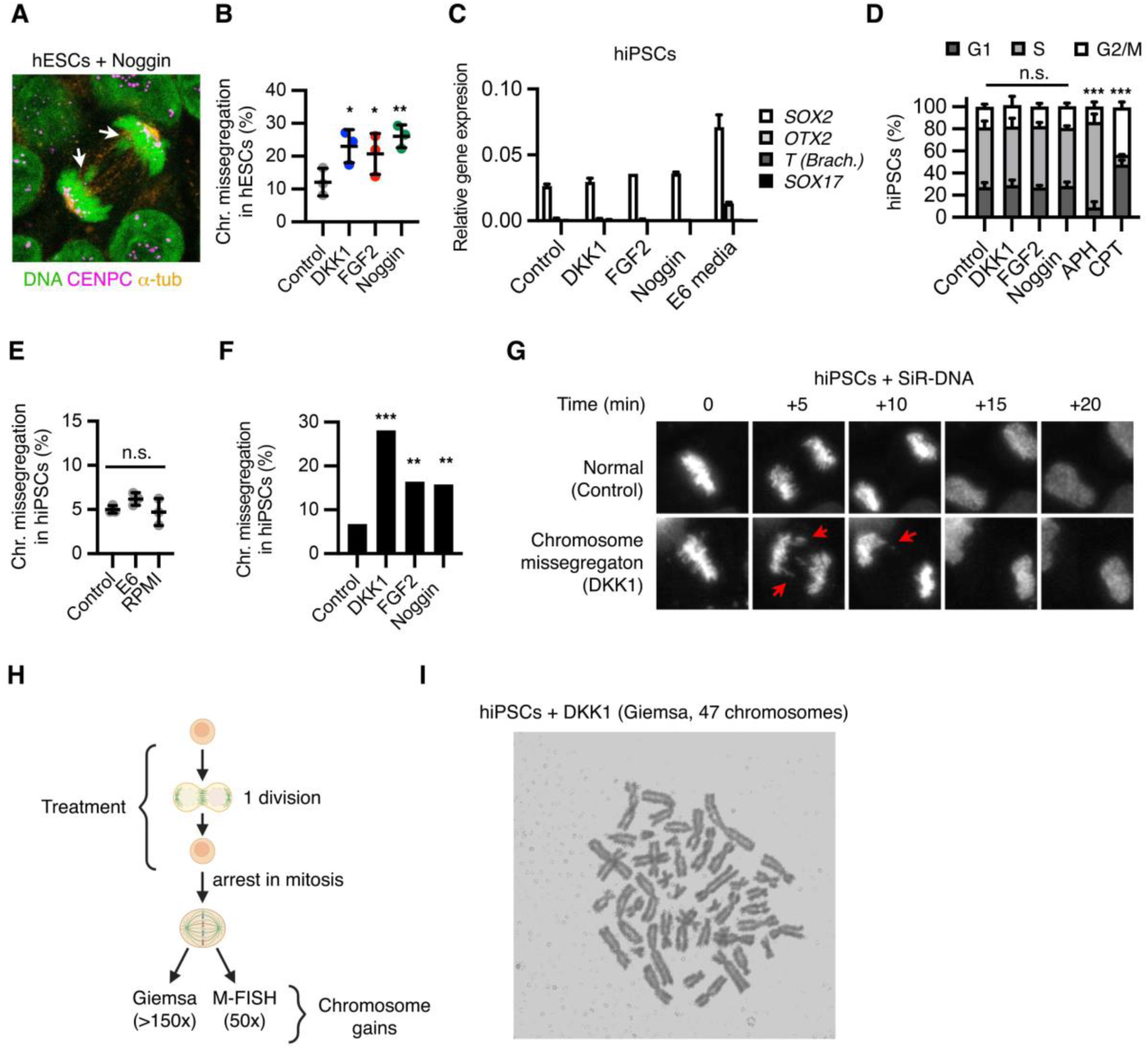
WNT, BMP and FGF regulate chromosome segregation fidelity in pluripotent stem cells. **A,B**, Chromosome segregation analyses in human embryonic stem cells (hESCs) upon treatment with the indicated signals for 16 hours. **C**, Representative qRT-PCR analyses of hiPSCs treated for 16 hours as indicated. **D**, Cytometry analyses of hiPSCs treated as indicated. APH, aphidicolin; CPT, Camptothecin. **E**, Chromosome segregation analyses in hiPSCs upon culture for 16 hours in control medium (E8) or differentiating media (E6, RPMI). **F,G**, live cell imaging analyses of mitotic hiPSCs labelled with SiR-DNA and treated as indicated. Data are mean of 50 cells completing mitosis. **H**, Experimental setup for the karyotype analyses shown in Figure 2G. **I**, Example of Giemsa staining of a hiPSC treated with DKK1. *P*-values were calculated from one-way ANOVA analyses with Tukey correction, or Fisher’s exact test **(B, E, D,F)**, are indicated as *P < 0.05, **P < 0.01, ***P < 0.001, or n.s., not significant. Data are mean ± s.d. of 3 biological replicates

**Figure S3:**
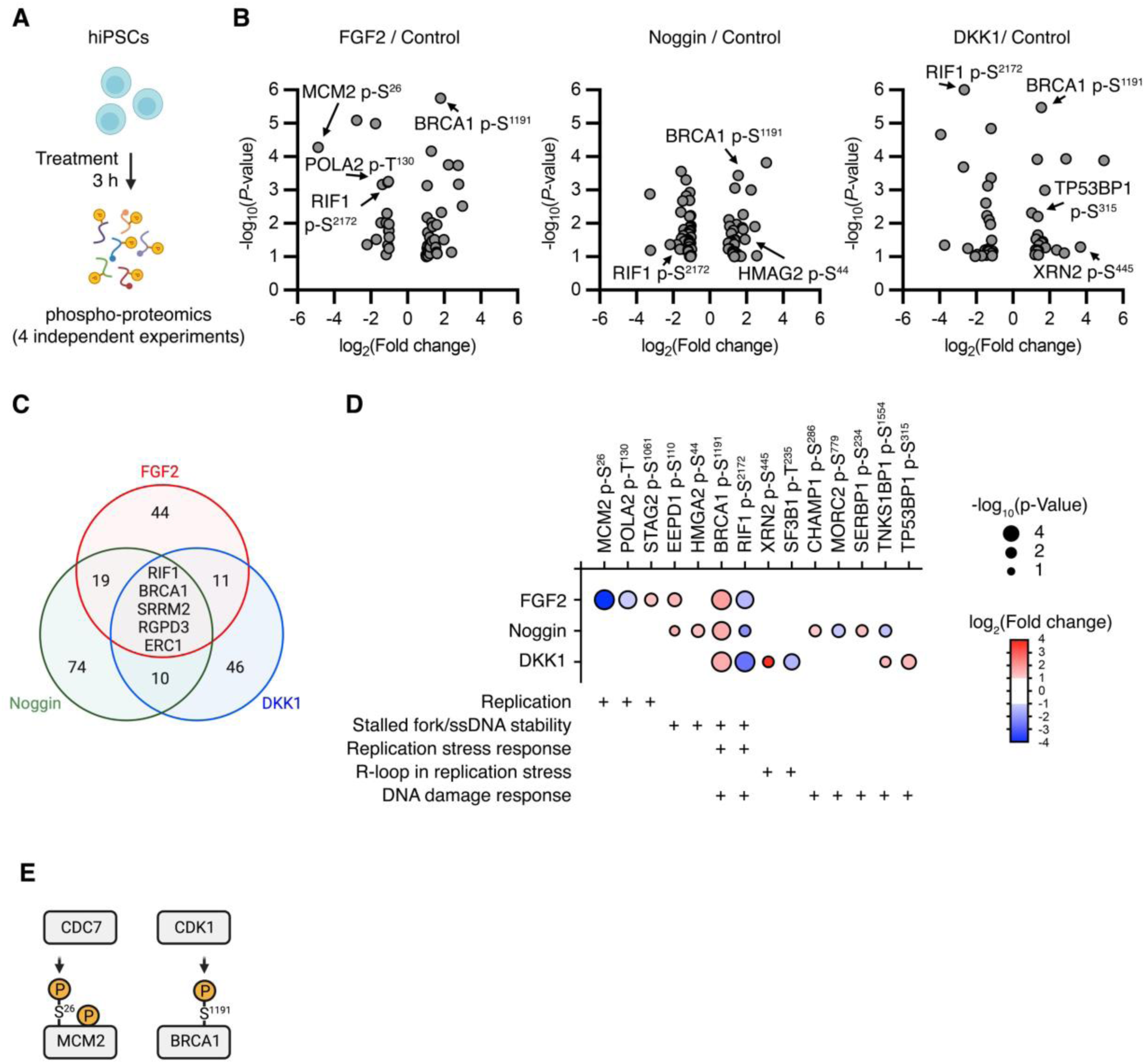
phospho-proteomics analyses upon WNT, BMP and FGF signalling perturbation. **A**, Schematic of the phospho-proteomics analysis. **B**, Differentially regulated phospho-peptides by DKK2, Noggin and FGF2 in hiPSCs after 3 hours. Note that DNA replication and damage response proteins are among most significantly regulated factors by either treatment. **C**, Venn diagram of differentially regulated proteins in the phospho-proteome analysis. **D**, Differentially regulated factors from the phospho-proteomics experiment that are associated with DNA replication and/or damage response. The list was hand-curated by searching each differentially regulated proteins for known roles in those processes (See table 3). **E**, Schematic with two known kinase-target associations.

**Figure S4:**
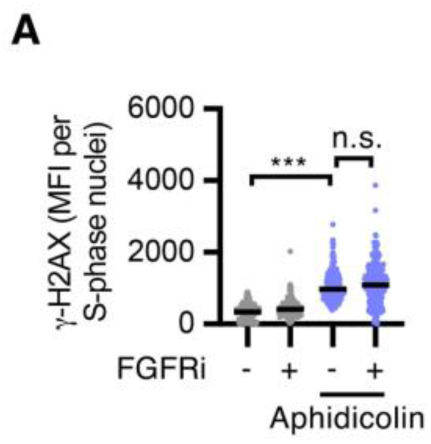
Epistatic analyses of FGF signalling in DNA replication stress. **A**, Accumulation of γ-H2AX foci in hiPSCs treated for 3 hours as indicated. Data are median fluorescence intensity (MFI) of > 200 EdU^+^ nuclei. *P*-values from one-way ANOVA analyses with Tukey correction are indicated as *P < 0.05, **P < 0.01, ***P < 0.001, or n.s., not significant.

**Figure S5:**
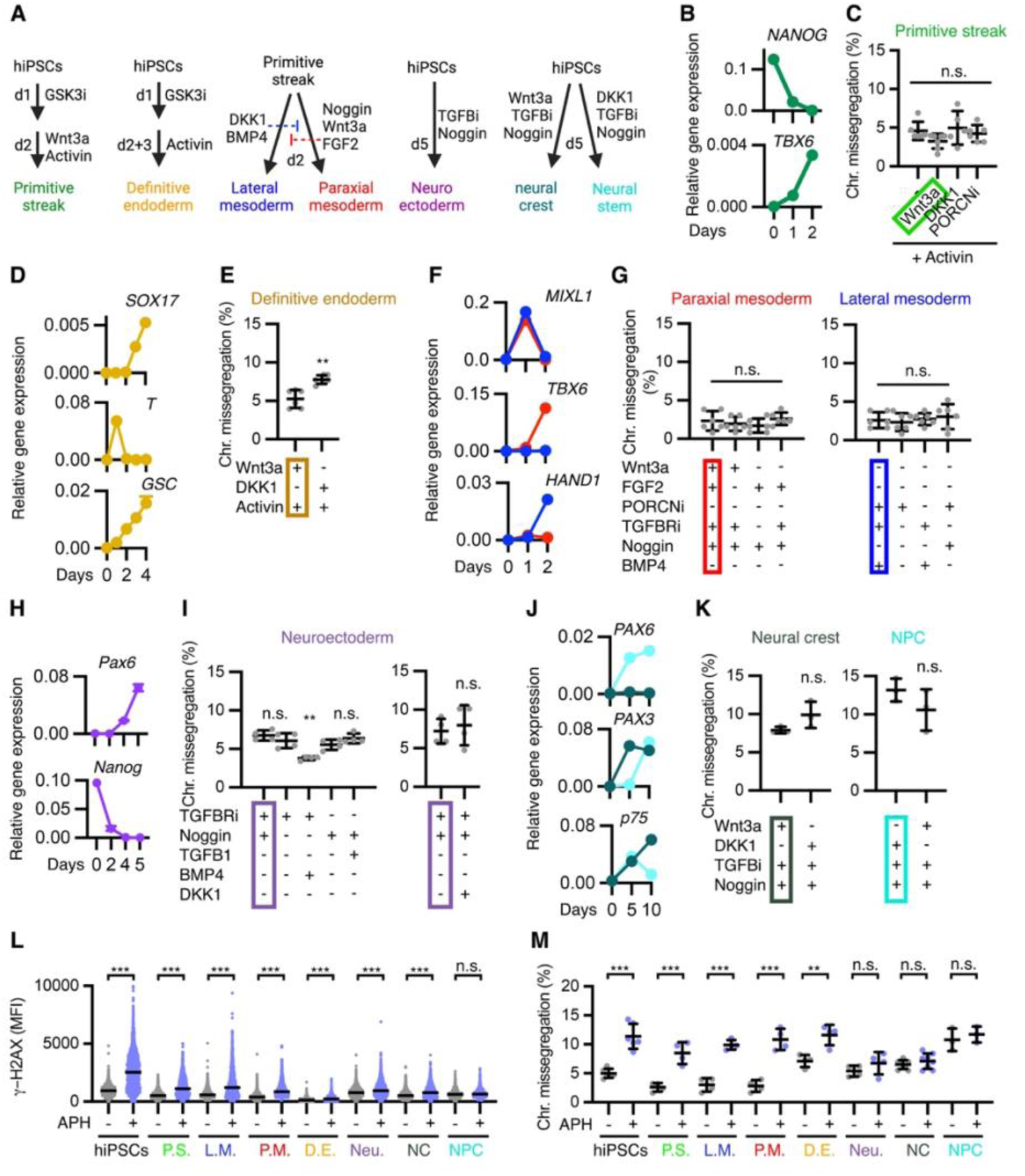
WNT, BMP and FGF display limited functions in chromosome segregation fidelity after lineage specification. **A**, Summary of the treatments and days (d) required to generate different lineages from hiPSCs. **B-K**, Cell lineage specification experiments performed as indicated in **(A)**. In **(B, D, F, H, J)** cells were harvested at the indicated differentiation days and analysed by qRT-PCR for lineage-specific markers. In **(C, E, G, I, K)** differentiating media was modified as indicated during the last 16 h and chromosome segregation was analysed by immunofluorescence. Data are mean ± s.d. of biological replicates with >100 cells per condition**. L, M**, Impact of aphidicolin (APH)-induced DNA replication stress in DNA damage and chromosome missegregation in the indicated lineages, generated as shown in **A)**. In **(L)**, data are median fluorescence intensity (MFI) of γ-H2AX of > 300 nuclei. In **(M)**, data are mean ± s.d. of biological replicates with > 100 cells per condition. Primitive streak-like (P.S.), lateral mesoderm (L.M.) paraxial mesoderm (P.M.), definitive endoderm (D.E.), neuroectoderm (Neu), neural crest (NC) and neural progenitor cells (NPC) are shown. *P*-values from one-way ANOVA analyses with Tukey correction are indicated as *P < 0.05, **P < 0.01, ***P < 0.001, or n.s., not significant. Colour boxes indicate the standard conditions for differentiation into the indicated lineages.

**Figure S6:**
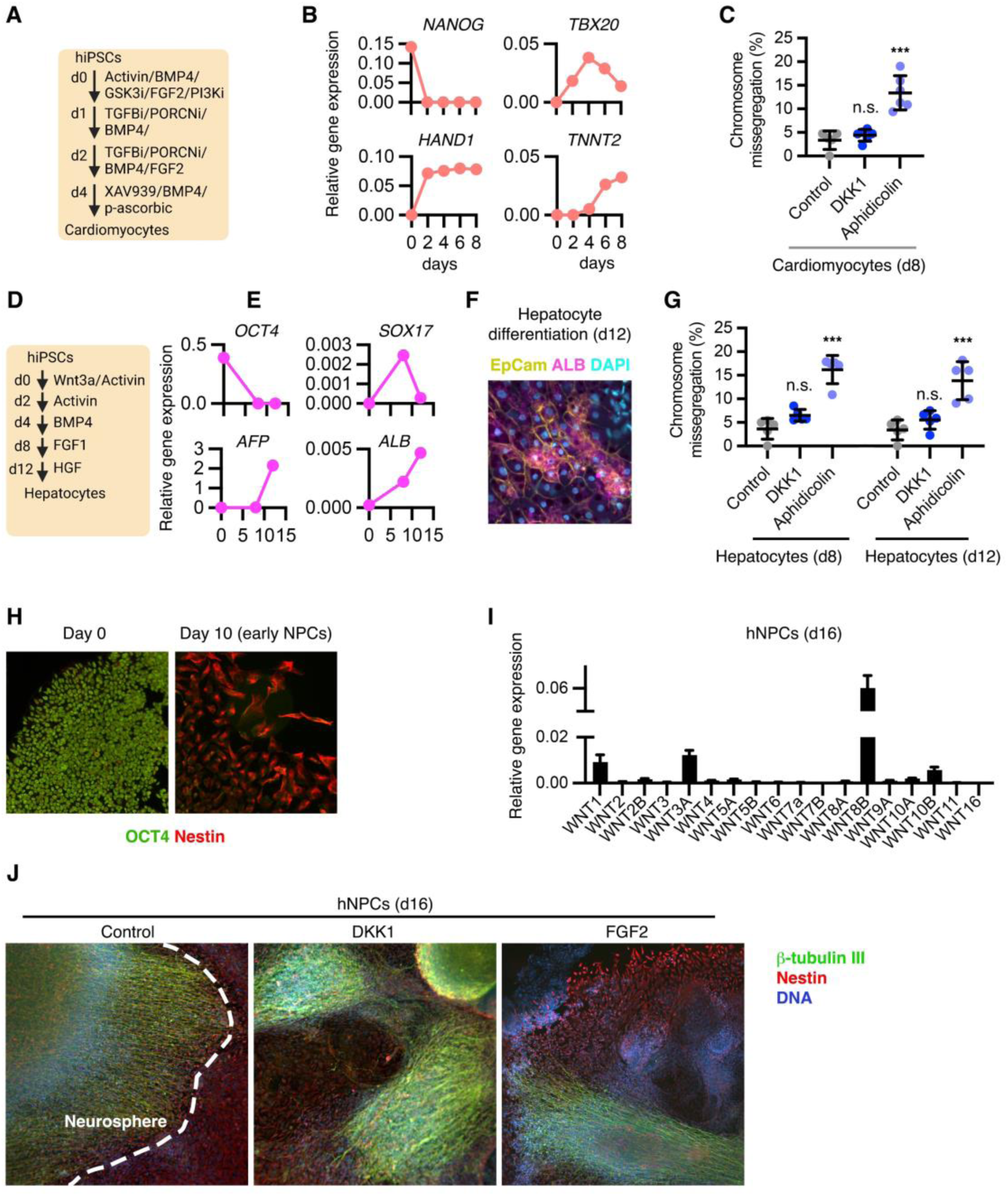
*In vitro* differentiation of hiPSCs into hepatocyte-, cardiomyocyte-like cells and neural progenitors. **A**, Summary of the treatments and days required to differentiate hiPSCs into cardiomyocyte-like cells. **B**, Representative qRT-PCR analyses of hiPSCs undergoing differentiation into cardiomyocytes. **C**, Chromosome segregation analyses of *in vitro* generated cardiomyocyte-like cells treated for 16 hours as indicated. **D**, Summary of the treatments and days required to differentiate hiPSCs into hepatocyte-like cells. **E**, Representative qRT-PCR analyses of hiPSCs undergoing differentiation into hepatocytes. **F**, Immunofluorescence analysis of immature hepatotcyte-like cells at day 20. **G**, Chromosome segregation analyses of *in vitro* generated hepatocyte-like cells treated for 16 hours as indicated. **H**, Immunofluorescence analyses of hiPSCs (Day 0) and expanding hiNPCs (Day 10). **I**, qRT-PCR analyses of WNT ligands in hiNPCs at day 16 of differentiation. **I**, Immunofluorescence analyses of hiNPCs at day 16 of differentiation, treated for 16 hours as indicated.

**Figure S7:**
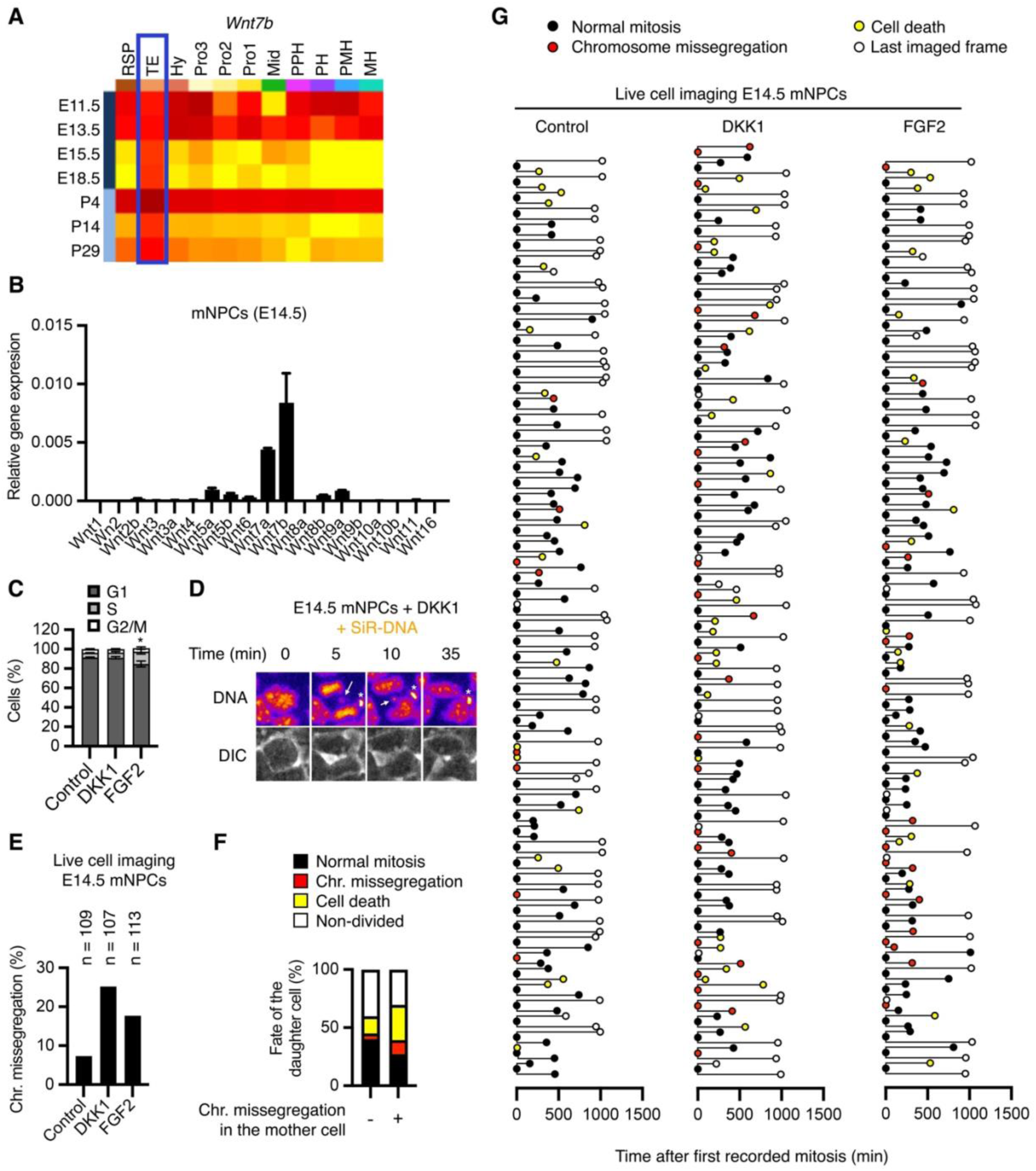
Signalling and chromosome segregation analyses in mouse NPCs. **A**, Expression profile of *Wnt7b* in the developing nervous system obtained from the Allen Developing Mouse Brain Atlas (http://developingmouse.brain-map.org/). RSP: rostral secondary prosencephalon; TE: telencephalon (highlighted); Hy: peduncular (caudal) hypothalamus; Pro3: prosomere 3; Pro2: prosomere 2; Pro1: prosomere 1; Mid: midbrain; PPH: prepontine hindbrain; PH: pontine hindbrain; PMH: pontomedullary hindbrain; MH: medullary hindbrain (medulla). **B**, Representative qRT-PCR analyses of WNT ligands in NPCs isolated from E14.5 mouse embryos. **C**, Cell cycle profiles of mNPCs isolated from E14.5 embryos and treated for 16h as indicated. Data are mean ± s.d. of 3 independent FACS experiments. *P*-values from one-way ANOVA analyses with Tukey correction are indicated as *P < 0.05. **D-G**, Representative live cell imaging analyses of NPCs isolated from E14.5 mouse embryos and stained with SiR-DNA. In (**D**), an exemplary mitosis of NPCs treated with DKK1 is shown. The arrow indicates a lagging chromosome and the asterisk marks an artefact. In (**F, G**), the fate of daughter cells following a normal or chromosome missegregation division in the mother cell was tracked during the recording. In (**F**), all tracked divisions (n = 350) from control-, DKK1– and FGF2-treated mother cells are shown. Note that chromosome missegregation in mother cells do not change the proportion of daughter cells committed to division (Normal mitosis + Chr. Missegregation). In (**G**), single tracks for the daughter cells of divisions in each individual treatment are shows.

## Supplemental Methods

### Cell culture

The human induced pluripotent stem cells (hiPSCs) were a gift from Kyung-Min Noh (EMBL). Cells were seeded in wells coated with Vitronectin 1h at 37°C (VTN 1:100 diluted in PBS), and grew in hiPSC culture medium. In detail, cells were cultured in Essential E8 medium (Thermo Scientific) supplemented with Penicillin/Streptomycin. We added Revitacell Supplement (Thermo Scientific) for the first 24 h after plating. Media were changed every day, and cells were split every 3 to 4 days using Versene (Gibco). All the experiments were carried out with hiPSCs between passage 10 and 20. For immunofluorescence experiments, 50.000 cells were seeded per each 12-well.

The mouse feeder-free embryonic stem cell line Sox1-GFP was a gift from A. Smith (University of Cambridge), and the line E14Tg2a a gift from C. Niehrs (DKFZ). Cells were cultured in DMEM+15% Panserum (PANSERA) supplemented with Leukemia Inhibitor Factor (LIF) and Penicillin/Streptomycin. Prior to seeding, well plates were coated with Gelatin 0.1% diluted in PBS for 10 min at 37C. Routine passage was carried out every 2-3 days, after seeding 100.000 cells/well in 6-well plates. For immunofluorescence experiments, 50.000 cells were seeded per each 12-well.

H9 hESCs were cultured in 6 well plates pre-coated with 1.6% Growth Factor Reduced Matrigel (356231, Corning) diluted in DMEM F12 (21331-020, Thermo Fisher Scientific) for 30 minutes at 37 °C in mTeSR Plus (100-0276, StemCell Technologies). hESCs were routinely passaged with EDTA (produced in-house) at a splitting ratio of 1 to 5 every 3 days, with media change every day.

Human WNT3A, DKK1-FLAG (DKK1) and RSPO3-ΔC-AP (RSPO3) ^90^ conditioned media were generated in-house using the corresponding basal media of interest and tested regularly in WNT reporter assays, as previously indicated ^57, 91^.

### Cell lineage specification experiments

Embryoid bodies (EBs) were generated from hiPSCs to validate the conserved functions of morphogens in human gastrulation. After 3-4 days in culture, hiPSCs were dissociated using 0.5mM EDTA in PBS, washed in PBS and reaggregated in E6 basal differentiation medium, with the corresponding anteriorising or posteriorising signals (as described in Figure S1) and supplemented with 1:2000 ROCK inhibitor (Y-27632). Cells were counted using a Neubauer chamber, and 1000 cells were seeded per well in an ultra-low-adherence 96-well plate (CellStar, 650970). 24h after aggregation, cells were harvested for RNA extraction and qPCR analysis (Figure S1).

Primitive streak, mesoderm and endoderm lineages were generated as previously described^27^. Briefly, we plated hiPSCs in low confluency (1:25) for 2 days in hiPSC culture medium prior differentiation. During the differentiation procedures, we grew cells in A-RPMI media supplemented with GlutaMax and Penicillin/Streptomycin. To generate primitive streak-like cells, we treated hiPSCs with 5 μM CHIR99021 for 1 day, followed by overnight treatment with WNT3A-conditioned medium (1:5) and 50 ng/mL Activin A. For paraxial mesoderm, we treated hiPSCs with 4 μM CHIR99021, 30 ng/mL Activin A, 20 ng/mL FGF2 and 100 nM PIK90 for 1 day, followed by treatment with 3 μM CHIR99021, 1μM A830, 250 nM LDN and 20 ng/mL FGF2 for another day. For lateral mesoderm, we treated hiPSCs with 6 μM CHIR99021, 30 ng/mL Activin A, 40 ng/mL BMP4, 20 ng/mL FGF2 and 100 nM PIK90 (PI3Ki), followed by 1-day treatment with 1 μM A8301 (TGFBRi), 30 ng/mL BMP4 and 1 μM C59 (PORCNi).

Neuroectoderm, neural crest and neural stem cells were generated as previously described ^28^. Briefly, we plated hiPSCs in low confluency (1:35) for 1 day in hiPSC culture medium prior differentiation. During the differentiation, we cultured cells in E6 media supplemented with Penicillin/Streptomycin. To generate neuroectoderm, we grew hiPSCs in the presence of 200 ng/mL Noggin and 10 μM SB431542 (TGFBi) for 5 days, changing media on days 2 and 4 of the differentiation protocol. To generate neural crest-like cells, we treated hiPSCs for 5 days with 200 ng/mL Noggin, 20 μM SB431542, and 1:5 WNT3A conditioned medium, changing the media on day 2 and 4. To generate early neural progenitors, we treated hiPSCs for 5 days with 200 ng/mL Noggin, 20 μM SB431542, and 1:5 DKK1 conditioned medium, changing the media on day 2 and day 4.

Hepatocyte-like cells were generated from hiPSCs as described before ^92^. Briefly, we plated hiPSCs in low confluency (1:35) for 2 days in hiPSC culture medium prior differentiation. During the differentiation procedures, we grew cells in A-RPMI media supplemented with GlutaMax and Penicillin/Streptomycin. To generate hepatocyte-like cells, we treated hiPSCs with 1:5 Wnt3a, 50 ng/mL Activin A, and 0.6% DMSO for 2 days, followed by 50 ng/mL Activin A, 0.6% DMSO for another 2 days to generate definitive endoderm. To further differentiate into hepatocyte-like cells, we treated the cells with 50 ng/mL BMP4 and 0.6% DMSO for 2 days, followed by 2 days with 20 ng/mL FGF1 and 0.6% DMSO. To generate more mature hepatocyte-like cells, cells were treated with 20 ng/mL HGF supplemented with 2% DMSO until day 20 of differentiation. Before changing the composition of the growth medium, cells were washed with DMEM/F12. Medium was changed every 2^nd^ day.

Cardiomyocyte-like cells were generated as previously described ^27^. Briefly, we plated hiPSCs in low confluency (1:25) for 2 days in hiPSC culture medium prior differentiation. During the differentiation procedures, we grew cells in A-RPMI media supplemented with GlutaMax and Penicillin/Streptomycin. To generate cardiomyocyte-like cells, we treated hiPSCs with 30 ng/mL, 40 ng/mL BMP4, 6µM CHIR99021, 20 ng/mL FGF2, and 100 nM PIK90 for 1 day, followed by another day culture with 1µM A8301, 30ng/mL BMP4, 1µM + C59 to generate lateral mesoderm. To further differentiate into cardiac-mesoderm, we treated the cells with 1µM A8301, 30ng/mL BMP4, 1µM C59, and 20 ng/mL FGF2 for 2 days, followed by 30 ng/mL BMP4, 1 µM XAV939, and 200 µg/mL Phospho-ascorbic-acid to reach cardiomyocyte-like cells. Before changing the composition of the growth medium, cells were washed with DMEM/F12. Medium was changed every 2^nd^ day.

Human induced neural progenitors (hiNPCs) were generated using previous protocols with few modifications^66^. In detail, hiPSCs were cultured in hiPSC culture medium for 4 days until reaching 70-80% confluence. Cells were dissociated using Dispase (5 min at 37C), washed with PBS, and resuspended in T75 flasks medium consisting 50% E8 Medium + 50% Neural Induction Medium (NIM, DMEM/F12 (Gibco) supplemented with 1× N2 (Gibco), 1× non-essential aminoacids and 1× Penicilin/Streptomycin) in order to aggregate in non-adherent conditions (Day 0). Media was changed on Day 1 (50% E8 Medium + 50% NIM), prior culture from Day 2-7 in daily changed 100% NIM media by slowly centrifuging the cell aggregates (embryoid bodies) at 200 RPMs. On Day 7, 12-well plates with glass coverslips were coated using poly-D-Ornithine (15 μg/mL) 3h at 37°C. Embryoid bodies were collected and seeded at a confluence of around 15-20 EBs/12-well in NIM media. Media was changed every two days, and cells were harvested on Day 10 (early NPCs) and 16 (late/mature NPCs).

In Figure 6 and Figure S6J, signal molecules were modified during the last 16 h of treatment as indicated in each individual panel.

### Protein and small compound treatments

Where indicated, cells were treated for 16 h or 3 h (Fig. 2C-J, Fig. 4E-G, and Fig. S1) with 1:3 human DKK1 conditioned medium, 1:3 human WNT3A conditioned medium, 40 ng/mL recombinant human FGF2 (R&D), 5 ng/mL recombinant human TGFB1 (Peprotech), 200 ng/mL recombinant human Noggin (R&D), 100 ng/mL recombinant human EGF (R&D), 100 ng/mL recombinant human BMP4 (Peprotech), 50-200 nM aphidicolin (Sigma-Aldricht), 3 μM AZD0156 (ATMi) (Selleckchem), 2.5 μM CPT (Camptothecin, Selleckchem), 3 mM hydroxyurea (HU), 3 μM CHIR99021 (GSK3i) (Selleckchem), 20 μM nucleoside mix (dNs) (SCBT), 10 nM taxol (Sigma), 0.1 μM FGFRi (Merck), or 10 μM LGK-974 (Selleckchem). A complete list of morphogens, growth factors, and small compounds as well as the concentration they were used in Figure 1B is shown in table 1.

### Chromosome segregation and DNA damage analyses

For chromosome segregation analyses, cells were treated for 16 h, as indicated. For DNA damage analyses (γ-H2AX) cells were supplemented with EdU for 30 min prior treatment to visualize S-phase cells, and after treated for 3 h prior collection. Culture cells were fixed in 2-4% PFA for 10-15 min, permeabilised with 0.5% Triton X100 in PBS (PBST) for 10 min, followed by a blocking step for 20 min and overnight incubation with primary antibodies in 2% horse serum in 0.25% PBST. We used 1:250 guinea pig anti-CENPC antibody (MBL International Corporation, USA, cat no PD030) to stain kinetochores (Chromosome segregation), or 1:250 mouse anti-phospho-H2AX (Ser139) (γ-H2AX) antibody (Millipore, clone JBW301) to detect DSBs, or 1:250 mouse anti-phospho-RPA32/RPA2 (Ser4 + 8) (pRPA) antibody to detect single strand DNA breaks; and the secondary antibodies 1:500 anti-guinea pig Cy3 (Millipore, AP308P), 1:500 anti-mouse Alexa488 (ThermoFisher A21202), supplemented with 1 μg/ml DAPI. In Figure 2G, incorporated EdU was subjected to click-it reaction (ThermoFisher, C10337), as indicated by the manufacturer.

H9 hESCs were platted at a ratio of 1:15 in 12 well plates on the top of 15mm coverslips pre-sterilized in ethanol 100%, washed in sterile ddH2O and coated with 1.6% Growth Factor Reduced Matrigel diluted in DMEM F12. Two days after plating, cells were treated with 250 ng/mL of DKK1; 40 ng/mL of FGF2, or 200 ng/mL of Noggin for 16 hours. Cells were fixed using 4% paraformaldehyde (PFA) (15710, Electron Microscopy Sciences) diluted in PBS for 20 minutes at room temperature and then washed three times with PBS – 0.1% Tween. The permeabilisation step was performed in PBS containing 0.3% Triton X-100 and 0.1 M glycine for 30 minutes at room temperature. Samples were incubated with primary antibodies (CENPC – MBL, PD030 and α-tubulin – Sigma, T9026) diluted at 1:250 in blocking solution (3% BSA and 0.1% Tween in PBS) overnight at 4 °C. The day after, samples were washed with PBS – 0.1% Tween three times, followed by incubation with secondary antibodies (donkey anti-guinea pig 647, AP193SA6 Millipore, and donkey anti-mouse 488, A-21202 ThermoFisher Scientific), diluted at 1:500 in blocking solution for 2 hours at room temperature.

To quantify cells exhibiting chromosome missegregation, we analysed 100-200 anaphases (hiPSCs, mESCs) and 50-100 anaphases (mNPCs, hiNPCs, lineages) in each biological replicate using either a Nikon Eclipse Ti using a 60× objective with oil immersion; or an inverted SP8 confocal microscope (Leica Microsystems) with a Leica 63×/1.4NA Oil objective and an upright Olympus VS200 slide scanning microscope with a 40×/0.95NA Air objective (hESCs). In mESCs, hESCs, hiPSCs and their derived lineages, chromosomes clearly separated (Lagging) from the bulk of segregated DNA chromatids were considered as chromosome missegregation. In mouse NPCs, which display lower distance between the two masses of separating chromosomes, we quantified chromosomes clearly separated from or bridging between the bulk of segregated DNA, as previously shown (Fig. 5G, ^10^; Fig.4A, D, ^20^). Basal levels of chromosome missegregation in hiPSCs and mouse NPCS (E14.5) were in accordance to previous estimates ^10, 37^.

To characterise DNA damage, EdU (Fig. 2G and Fig. S3B) or DAPI stained nuclei were automatically selected, and the median fluorescence intensity (MFI) calculated, using ImageJ Fiji 2.0.0-rc-69/1.52p, after background subtraction.

### DNA combing experiments

Single molecule DNA combing assays in hiPSCs and hiNPCs were performed to determine replication fork progression speed and inter-origin firing distances (OFD). Cells were treated with proteins or small molecule compounds 3 hours as indicated before harvesting. To label the replication forks, cells were incubated with two 30 min consecutive pulses of 100 μM 5-Chloro-2ʹ-deoxyuridine (CldU; Sigma-Aldrich) and, 100 μM 5-iodo-2ʹ-deoxyuridine (IdU; Sigma-Aldrich), respectively. Cells were harvested and processed using the FiberPrep DNA extraction kit (Genomic Vision, France), as indicated by the manufacturer. Isolated DNA was immobilized on vinylsilane engraved coverslips (Genomic Vision, France) using the Molecular Combing System (Genomic Vision, France) at 2 kb/μm. Coverslips were stained with 1:10 anti-BrdU/CldU (BU1/75 (ICR1), Abcam, ab6326), 1:10 anti-BrdU/IdU (B44, BD Biosciences, 347580), 1:5 anti-ssDNA (IBL, 18731), and secondary conjugated antibodies to Cy5 (1:25, Abcam, UK, cat no ab6565), Cy3.5 (1:25, Abcam, UK, ab6946), and BV480 (1:25, BD Biosciences, 564877). Images were acquired using a Nikon CREST microscope using a 60× objective with oil immersion and autofocus (ssDNA). Using NIS Elements software, we stitched together 5 wide-field images along the longitudinal axis of the combed DNA. We analysed at least 100 labelled unidirectional DNA tracks per sample to determine replication fork progression rates. To determine inter-origin firing distances, the distance between two neighbouring origins on the same DNA strand was measured for at least 45 origin pairs per condition.

### Karyotype analyses

Cells were treated for 16 h, as indicated, followed by 16 h arrest using 100 ng/mL nocodazole.

Giemsa staining and karyotype analysis was performed as described previously ^23^. Briefly, cells were pelleted, washed with PBS, and incubated in hypotonic medium (40% DMEM-F12, 60% H2O) at RT for 15 min. Cells were fixed in Carnoy’s solution (methanol:acetic acid = 3:1). Chromosomes spread onto glass slides and stained with Giemsa solution, and imaged using a Nikon Eclipse Ti with a 60× oil immersion objective.

M-FISH was performed as previously described ^93^. Briefly, seven pools of flow-sorted human whole chromosome painting probes were amplified and combinatorial labelled using DEAC-, FITC-, Cy3, TexasRed, and Cy5-conjugated nucleotides and biotin-dUTP and digoxigenin-dUTP, respectively, by degenerative oligonucleotide primed (DOP) PCR. Prior hybridisation, metaphase spreads fixed on glass slides were digested with pepsin (0.5 mg/ml; Sigma) in 0.2N HCL (Roth) for 10 min at 37°C, washed in PBS, post-fixed in 1% formaldehyde, dehydrated with a degraded ethanol series and air dried. Slides were denatured in 70% formamide/1× SSC for 2 min at 72°C. Hybridization mixture containing combinatorial labelled chromosome painting probes, an excess of unlabelled cot1 DNA in 50% formamide, 2× SSC, and 15% dextran sulphate were denatured for 7 min at 75°C, pre-annealed for 20 min at 37°C, and hybridized to the denaturated metaphase preparations. After 48 h incubation at 37°C slides were washed 3 times at room temperature in 2× SSC for 5 min, followed in 0.2× SSC/0.2% Tween-20 at 56°C 2 times for 7 min. For indirect labelled probes, an immunofluorescence detection was carried out. Therefore, biotinylated probes were visualized using three layers of antibodies: streptavidin Alexa Fluor 750 conjugate (Invitrogen), biotinylated goat anti avidin (Vector) followed by a second streptavidin Alexa Fluor 750 conjugate (Invitrogen). Digoxigenin labelled probes were visualized using two layers of antibodies: rabbit anti-digoxin (Sigma) followed by goat anti-rabbit IgG Cy5.5 (Linaris). Slides were washed in between 3 times in 4× SSC/0.2% Tween-20 for 5 min, counterstained with 4.6-diamidino-2-phenylindole (DAPI) and covered with anti-fade solution. Images of metaphase spreads were taken for each fluorochrome using highly specific filter sets (Chroma technology, Brattleboro, VT) on a DM RXA epifluorescence microscope (Leica Microsystems, Bensheim, Germany) equipped with a Sensys CCD camera (Photometrics, Tucson, AZ). Camera and microscope were controlled by the Leica Q-FISH software and images were processed on the basis of the Leica MCK software and presented as multicolour karyograms (Leica Microsystems Imaging solutions).

### Single cell RNA-sequencing

For single cell sequencing analyses, hiPSCs were treated for 16 h with 1:3 DKK1 conditioned medium, 40 ng/mL recombinant FGF2 (R&D), or 200 ng/mL recombinant Noggin (R&D). Live hiPSCs were stained with DAPI to exclude death cells, Annexin V to exclude pre-apoptotic cells, and DRAQ-5 to sort G1 cells. Cells were FACS sorted in 5μL lysis buffer (Quiagen), and the RNA separated using Biotin-dT30-bound streptavidin beads (ThermoFischer, 18064014). After cDNA synthesis and amplification using SuperScript III Reverse transcriptase kit (Thermo), samples were cleaned-up with 0.6× SPRI beads and tagmented with homemade Tn5 at 55°C for 3 min. We performed a final PCR amplification of 12 cycles using KAPA HiFi kit (Roche) with unique pairs of i5/ i7 adapter index primer, proceeded with 0.8× SPRI bead purification, pooled samples after DNA concentration quantification in Bioanalyzer, and performed a final 0.75× SPRI clean up. Samples were sequenced by NextSeq 2000 (Illumina) and with 75 paired-end (PE) reads. Read alignments and the count tables of mapped read per gene were obtained using STAR version 2.6.0a with GRCh38 human reference genome an its gene model (GRCh38.84). For differential gene expression analysis, the R package Seurat v 4.1.1 ^94^ was applied. We filtered out genes that were detected in less than 3 cells and selected high quality cells that had less than 25% mitochondrial counts and more than 4,000 and less than 12,000 detected genes. After normalizing the data for differences in library size, the ‘FindMarkers’ function with the ROC test was used to determine differentially expressed genes (DEGs) between the different treatment conditions. We selected DEGs with power > 0.25 for the subsequent Gene Ontology (GO) analysis (https://david.ncifcrf.gov/tools.jsp, 07/2022).

### qRT-PCR and FACS analyses

RNA was extracted and purified using Quiagen RNAeasy Plus column kit, according to the manufacturer’s instructions. The cDNA was produced with SensiFAST cDNA Synthesis kit (Bioline) starting from 300 ng to 1 μg mRNA. Real-time quantitative PCR reactions from 8,3 ng of cDNA were set up in technical triplicate using the SensiFAST SYBR Hi-ROX kit (Bioline) on a StepOne Plus qPCR machine (ThermoScientific). The sequences of the oligonucleotides used in this study are provided on request. Expression levels were normalized to PCR amplification with primers for *GAPDH*.

Cell cycle profiles were performed in hiPSCs treated as indicated for 16 hours with the selected factors. Prior harvesting, cells were pulsed 1 hour using BrdU (Sigma). Cells were collected in ice cold PBS, fixed in ice cold ethanol (final concentration 70%) and stained using an anti-BrdU antibody and propidium iodide as described before ^91^.

### MS sample preparation and analysis

hiPSCs were seeded in 10 cm dishes in E8 media and treated with control, DKK1, Noggin or FGF2 (See table 1) for 3 h before harvesting. Four independent experiments were processed for phospho-proteome analysis was performed as described previously ^57^. In brief, proteins were extracted by full lysis of cells, digested, enriched for phospho-peptides with TiO_2_ beads, and analysed by mass spectrometry. For the downstream analysis, only peptides with fold changes > 2 and p-values < 0.1 for the 4 experiments were considered differentially regulated. Each of the factors were manually analysed in STRING, PhosphoSitePlus, and other databases for i) their role in DNA replication or/and damage, ii) known kinases modulating the identified phospho-sites.

### Live cell imaging

For chromosome missegregation tracking, hiPSCs and mNPCs were cultured as described before in μ-Slide 8 Well chambers, and incubated with 500 nM SiR-DNA (Spirochrome AG, SC007) 1 h prior to and during the experiment. In preliminary analyses, we validated that this concentration of SiR-DNA did not induce mitotic delay or phenotypes. Treatments with selected factors were applied overnight before starting the imaging. Live cell imaging was performed using an automated Nikon Eclipse Ti2 inverted microscope equipped with a 40× water immersion objective (Nikon Apo LWD, NA 1.15) and a NEO sCMOS camera (Andor). Multipoint acquisition was controlled by NIS Elements 5.1 software. 5 z-stacks with 2-µm interval were recorded every 5 min for up to 5 h in a preheated chamber (STXG-WSKM, Tokai Hit) at 37 °C and 5% CO_2_. Images were analysed using ImageJ 2.0.0 software.

For microtubule dynamics analyses, cells were seeded onto 8-well glass-bottom imaging chambers (Ibidi) and transfected the next day with pEGFP-EB3 using Lipofectamine 3000. Cells were treated for 16 h, as indicated. Before imaging, cells were treated for 30 min with 2 μM Dimethylenastron to induce monopolar spindles, as previously described ^23^. Cells were live-imaged using a 100× 1.45 NA oil objective, in 4× 0.4 µm Z-optical sections with additional 1.5× magnification switch, every 2 s for over 30 s, in a humid chamber with 37°C, 5% CO_2_.

### Western Blotting

For Western blotting, cells were lysed in full lysis buffer (50 mM Tris⋅HCl, pH 7.5, 150 mM NaCl, 1% Nonidet P-40, 0.05% SDS, 1 mM β-mercaptoethanol, 2 mM EDTA, 1× protease phosphatase inhibitor mixture [Thermo Scientific]). The cleared lysates were mixed with 4× NuPAGE LDS sample buffer (Thermo Scientific) containing 50 mM DTT, resolved on 10% NuPAGE gels and transferred to nitrocellulose membranes. For Western blot experiments the following antibodies were used: rabbit anti–GAPDH (14C10 mAb, Cell Signalling, #2118), rabbit anti-phospho-Chk2 Thr68 (C13C1, Cell Signalling #2197) and rabbit-anti-phospho-Chk1 Ser345 (133D3, Cell Signalling, #2348).

### Mouse studies

E13.5 C57BL/6N wild type embryos were injected *in utero* either with 5 ng/μL DKK1 or with PBS+0,1% BSA control solution. In detail, pregnant mice were anesthetized with isoflurane, the uterine horns were exposed, and 1 μl of the solution was injected into the lateral ventricle of each embryo using glass micropipettes. Animals were sacrificed 16 hours later, and embryonic heads isolated in cold PBS, followed by fixation with 4% PFA for 3 days. Afterward, embryonic heads were cryoprotected in 30% sucrose solution and embedded in Tissue-Tek OCT. Embryonic coronal forebrain sections (18µm) were prepared using a Leica CM1950 cryo-microtome at the DKFZ Light Microscopy Core Facility. Cryosections were subjected to antigen retrieval using 1% sodium citrate, blocked in 0.1% PBST, and incubated overnight at 4°C with 1:250 anti-phospho-FGFR1, anti phospho-LRP6 S1490, anti-Nestin or/and anti-phospho-Histone 3.

Mouse neural progenitor cells were dissociated from the neocortex of E14.5 mouse embryo brains using the papain dissociation kit (LK003150, Worthington Biochemical Corporation) following the manufacturer instructions. Prior to seeding, 12-well plates with glass coverslips were coated overnight at 4°C using Poly-D-Ornithine (15 μg/mL), and rinse 3 times in PBS prior seeding. Dissociated NPCs were seeded at a 500,000 cells/cm^2^ density in medium consisting of: Neurobasal media supplemented with B27 (1×), Glutamax (1X) and Penicillin/Streptomycin. Media was changed daily and primary cultured cells were harvested after 48h for experiments.

### Ethics statement

Work with human embryonic stem cells (hESCs) was conducted at the MRC Laboratory of Molecular Biology (LMB) under an approval from the UK Stem Cell Bank Steering Committee, and in accordance with the regulations of the UK Code of Practice for the Use of Human Stem Cell lines. H9 hESCs were kindly provided by M. Lancaster (LMB) under an agreement with WiCell.

Time-mate pregnant C57BL/6N wild type mice for cortical isolations were purchased from Janvier. Animals had *ad libitum* access to food and water and were kept under a 12 h light – 12 h dark cycle. All animal experiments were approved by the local governing regional council under A.P. supervision. C57BL/6N wild type mice used for *in utero* injections were maintained and bred at the DKFZ central mouse facility. *In utero* injection experiments were approved by the local animal welfare committee *Regierungspräsidium Karlsruhe* following the guidelines from GV-SOLAS (AZ 35-9185.81/G-94/18 from J.A.). For terminal tissue harvesting procedures, pregnant mice were euthanized using cervical dislocation and embryos were decapitated, following the approved animal facility procedures.

### Statistical analyses

All the data and statistical significances were analysed using Prism 9 (GraphPad). Data are shown as mean with standard deviation of independent experiments or biological replicates of independent experiments, as indicated the figures; except for γ-H2AX analyses in Figures 2H, 4G, S3B, and S4L, where median values are represented. Where indicated, Student’s t-tests with two-tails (two groups) or ordinary one-way ANOVA analyses with Tukey correction (three or more groups) were calculated. In Figure 5A, fold changes and statistical analyses were calculated from the experiments shown in Figure S5. Significance of Gene Ontology groups from differentially expressed genes from Figure 3A was obtained from DAVID (https://david.ncifcrf.gov/tools.jsp, 07/2022). Significance is indicated as: *P < 0.05, **P < 0.01, ***P < 0.001, or n.s.: not significant.

## Supplementary Table legends

**Supplementary Table 1 (.xlsx file):** Technical details of the proteins and small compounds used in the chromosome segregation screen, as well as their potential role in gastrulation.

**Supplementary Table 2 (.xlsx file):** Differentially expressed genes (DEG) and Gene Ontology (GO) analyses in hiPSCs treated with DKK1, FGF2 or Noggin and analysed by single cell sequencing

